# Prospects and pitfalls of an intrusive model for the Châtelperronian stone tool industry during the Middle to Upper Palaeolithic transition in France and northern Spain

**DOI:** 10.1101/2023.07.14.549013

**Authors:** Igor Djakovic, Morgan Roussel, Marie Soressi

## Abstract

The Middle to Upper Palaeolithic transition in France and northern Spain reflects the transition from Neandertals to *Homo sapiens* and the emergence of novel cultural entities and standardised laminar technologies between ~50 and 40 thousand years ago. The Châtelperronian stone tool industry sits at the centre of this period, and is commonly considered as representing a geographically isolated archaeological entity produced by late Neandertals. However, debate as to the makers and origin of this industry has long persisted. Fuel has recently been thrown onto this discussion through the formulation of a hypothesis in which the Châtelperronian directly originates from the Northern Early Ahmarian industry of the Levant. This model proposes that the Châtelperronian is in fact indicative of a direct migration of a population of *Homo sapiens* from the Levant to France around 44-40 thousand years ago – potentially via the crossing (or series of crossings) of the Mediterranean Sea. Such a scenario would have significant implications for how we interpret this key portion of recent human evolutionary history. In this paper, we highlight some of the prospects and pitfalls of an intrusive origin model for the emergence of the Châtelperronian industry in western Europe - taking into account technological, chronological, geographic, and stratigraphic perspectives. To frame this discussion, we review the state of understanding on the Châtelperronian and provide a detailed, synthetic review of Châtelperronian lithic technology. Our review reinforces the distinctive and fully ‘Upper Palaeolithic’ character of this industry, and we subsequently suggest a few avenues of research which, in our opinion, may help shed progressively clearer light on the demographic and cultural processes operating during the Middle to Upper Palaeolithic transition in western Europe.

## 1. Introduction

The Middle to Upper Palaeolithic transition in France and northern Spain documents the progressive replacement of Neandertals by *Homo sapiens* in the region (Hublin, 2015). This period, taking place between ~55-40 thousand years ago, is from an archaeological perspective characterised by the emergence of distinctive stone tool industries with an increasing focus on standardised laminar lithic technologies (Teyssandier et al., 2010; Bordes and Teyssandier, 2011a; Metz et al., 2023; Slimak, 2023; Slimak et al., 2022). Namely, and in chronological order of appearance: the Neronian (limited to south-eastern France), the Châtelperronian, and the Protoaurignacian.

Of these industries, the Châtelperronian – which represents the first unambiguous departure from the Middle Palaeolithic in the region – is presently the only one which retains a Neandertal connection based on fossil-artefact associations (Leroi-Gourhan, 1958; Hublin et al., 1996; Bailey and Hublin, 2006a; Welker et al., 2016). This has played a key role in broader discussions concerning the relationship and/or interaction between Neandertals and *Homo sapiens* prior to the disappearance of the former circa 40 thousand years ago (e.g. Hublin et al., 1996; Mellars, 2010; Roussel et al., 2016). Perhaps most notably, following the emergence of the ‘acculturation’ model – which, in simple terms, supported the idea that the Châtelperronian is in some form the result of a diffusion of behaviours from migrant *Homo sapiens* onto local late Neandertals (e.g. Hublin et al., 1996; Mellars, 2010). The fact that the Châtelperronian has traditionally been thought to have emerged from a local Neandertal ‘substrate’ is of course at least partly due to the fact that analogous and contemporaneous stone tool assemblages have not been found in any other region of Europe.

Despite the fact that the makers and origin of the Châtelperronian have long been the subject of debate (Mellars et al., 2007; Bar-Yosef and Bordes, 2010; Higham et al., 2010; Bordes and Teyssandier, 2011a; Hublin et al., 2012; Gravina and Discamps, 2015; Roussel et al., 2016), there has been no explicit alternative hypothesis proposed as to the origin of this industry. Recently, this situation has changed. Slimak (2023) has put forward what we will refer to here as the ‘Three Waves’ model for the emergence of the Upper Palaeolithic in France and northern Spain. In simple terms, this model contends that the onset of the Upper Palaeolithic in this region - reflected by the Neronian, Châtelperronian, Protoaurignacian stone tool industry succession – is the result of three waves of AMH migration across the Mediterranean from the Levant to France and Spain between ~55 and 40 thousand years ago (*ibid.*). Such a scenario would have significant impacts on our current perspectives and interpretations for the onset of the Upper Palaeolithic and the disappearance of Neandertals in western Eurasia. Including, for example:

⮚ ***Homo sapiens* were the producers of the Châtelperronian industry and occupied large areas of France and northern Spain between ~45-40 kya, prior to the onset of the Protoaurignacian.** This would run contra to the model in which Neandertals are the sole authors of this industry – and would substantially alter our understanding of population demography for this ime period. Until recently, the Protoaurignacian was widely considered as a proxy for the first wide-scale *Homo sapiens* occupation in the region (Hublin, 2015). Another question which this would naturally raise is how to explain the presence of numerous Neandertal remains in the Châtelperronian levels of Groke du Renne (Maureille and Hublin, 2019).
⮚ **These groups of *Homo sapiens* were likely adept at sea-faring, making a crossing (or series of crossings) of the Mediterranean Sea from Lebanon to France around 44-40 thousand years ago.** Although clearly within the realm of viability given the arrival of *Homo sapiens* in Australia upwards of 65 thousand years ago (Clarkson et al., 2017), this is certainly not an empirically demonstrated model for the initial peopling of western Europe by *Homo sapiens*. Such a scenario would, additionally, open the door for alternative peopling scenarios for other coastal Mediterranean regions, for example Italy and Greece (Harvati et al., 2019) and the quasi-contemporaneous Uluzzian stone tool industry (Oxilia et al., 2022).
⮚ **If Mediterranean crossings did occur during the Middle to Upper Palaeolithic transition, Homo *sapiens* may have spread through Europe partly following a west-to-east (and not only east to west) axis.** This would be a significant shil in perspective with numerous implications. For example, the possibility of an earlier peopling of western Europe by *Homo sapiens* in comparison to central and/or eastern Europe. A common assumption within more traditional models is that this process must have essentially followed a land-based, east to west movement pakern (i.e. *Homo sapiens* should have a later occurrence in western Europe compared to eastern Europe) (e.g. Mellars, 2006; Hublin, 2015; Kadowaki et al., 2015). And in fact, chronological support for such a scenario has been consistently ambiguous despite increasingly higher quality chronological resolution for this period.
⮚ **There was likely a substanAally prolonged co-existence between Neandertals and *Homo sapiens* in France and northern Spain.** The widespread presence of *Homo sapiens* in this region between ~44 and 40 thousand years ago would substantially prolong the potential period of contact (Djakovic et al., 2022) between these human populations in this region. This is made further interesting by, for example, the recently published description of human remains from the site of La Coke de St Brelade (with a probably age of <48 kya) that indicate the presence of shared Neandertal/AMH traits in several permanent teeth (Compton et al., 2021).

Given the implications of such a scenario, in conjunction with the increasing lack of consensus amongst researchers as to the nature and origin of the Châtelperronian industry, it feels timely to present a critical review of this industry. In this paper, we highlight some of the prospects and pitfalls of an intrusive origin model for the emergence of this industry in western Europe - taking into account technological, chronological, geographic, and stratigraphic perspectives. To begin, we review the state of understanding on the Châtelperronian and provide a step-wise, synthetic review of Châtelperronian lithic technology. Our review reinforces the distinctive and fully ‘Upper Palaeolithic’ character of this industry and allows us to suggest a few avenues of research which, in our opinion, may help shed progressively clearer light on the demographic and cultural processes operating during the Middle to Upper Palaeolithic transition in western Europe.

## 2. Setting the scene

Viewed at the European scale, the Châtelperronian industry represents a notably coherent and geographically restricted archaeological entity (Soressi and Roussel, 2014). A little over 40 secure Châtelperronian sites have been recognized in France and northern Spain throughout an arch which stretches around 300 km wide, and fits closely to the western half of the Massif central (*ibid.*). This distribution stretches from Burgundy in north-central France, extends through south-west France to Cantabria in the west, and the south-eastern limit is marked by the Oriental Pyrenees. At present, no Châtelperronian assemblages have been identified east of the Rhône valley – although its northernmost limit has recently been extended to the site of Ormesson in the Paris basin (Bodu et al., 2017). Over the last decade, there has been substantial work aimed at refining the chronology of this industry and it is now widely accepted as belonging to a window of time between 44 and 40 kya cal BP (Hublin, 2015; Talamo et al., 2020; Djakovic et al., 2022) – and is found inter-stratified between Mousterian (below) and Protoaurignacian (above) layers when they occur at the same site (Soressi and Roussel, 2014).

The Châtelperronian was originally defined by H. Breuil after a lithic industry found at La Grotte des Fées at Châtelperron. Breuil emphasized the similarities between this assemblage and the Abri Audi type industry, later attributed to the Mousterian of Acheulean Tradition (MTA) – both of which were at the time considered as representing the first stage of the Upper Palaeolithic. This similarity was based partly on the presence of ‘backed knives’ which were found in both sets of assemblages. At the time, both industries were thought to have been produced by early groups of European AMHs. This perspective was shaken after the surprising discovery of a Neandertal skeleton in a Châtelperronian context at the site of Saint-Césaire (Lévêque and Vandermeersch, 1981). This proposed Neandertal-Châtelperronian association was later strengthened by the analysis of fossil discoveries from the Grotte du Renne, Arcy-sur-cure – the paleoanthropological evidence of which includes 29 isolated teeth, a temporal bone, and numerous fragmentary remains (Leroi-Gourhan, 1958; Hublin et al., 1996; Bailey and Hublin, 2006a).

The fossil-artefact associations at both sites have been the subject of considerable debate and critical re-evaluation (Bar-Yosef and Bordes, 2010; Higham et al., 2010; Caron et al., 2011; Hublin et al., 2012; Gravina et al., 2018). For Saint-Césaire in particular, it has recently been demonstrated that there exists no reliable association between the Neanderthal skeleton and the Châtelperronian/Mousterian component at this site (Gravina et al., 2018). The integrity of the Châtelperronian levels at Grotte du Renne has also been questioned, based largely on stratigraphic observations and the presence of a pronounced flake component within the assemblages (e.g. Bar-Yosef and Bordes, 2010; Connet, 2019). Despite this, in the current state of the art, a Neandertal authorship of the Châtelperronian remains the most empirically-supported model.

In terms of ‘origins’ for the Châtelperronian, the most commonly cited model posits a local origin of from the late Middle Palaeolithic of France – specifically, the MTA type B - with or without some form of ‘external’ influence from incoming AMHs producing Protoaurignacian technology (Soressi, 2005; Soressi and Roussel, 2014; Ruebens et al., 2015; Roussel et al., 2016a). As with the fossil-artefact associations, both the local origin model of the Châtelperronian and the external influence model have been the subject of important debate and critique (D’Errico et al., 1998; d’Errico et al., 2003; Gravina et al., 2005; Gravina et al., 2022; Mellars et al., 2007; Bar-Yosef and Bordes, 2010; Gravina and Discamps, 2015; Ruebens et al., 2015; Roussel et al., 2016a; Bodu et al., 2017). The consensus remains far from unanimous amongst researchers involved in the topic.

Recent chrono-cultural revisions of stratigraphic sequences in south-west France have created stratigraphic and chronological distance between the Châtelperronian and the MTA Type-B (Jaubert, 2011; Gravina and Discamps, 2015; Gravina et al., 2022). Specifically, it has been suggested that a Discoidal and possibly Levallois phase seem to separate the MTA and Châtelperronian at sites where they occur together (*ibid.*) – breaking the stratigraphic continuity which the MTA origin model has in part traditionally relied on. Furthermore, the Châtelperronian is now considered intrusive in the Iberian peninsula (Rios-Garaizar et al., 2022), and some have posited that a period of pronounced carnivore activity seems to have occurred between the final Middle Palaeolithic and Châtelperronian occupations in south-west France (Discamps, 2011).

The ‘Three Waves’ model has recently been put forward as an alternative scenario which would in some way resolve these ambiguities. In simple terms, this model proposes that the onset of the UP in this region - reflected by the Neronian, Châtelperronian, Protoaurignacian cultural succession – is the direct result of three distinct waves of AMH migration across the Mediterranean from the Levant to France (Slimak, 2023). The Châtelperronian would represent the second migration, and would be reflected in the Levant as what is called the Northern Early Ahmarian (NEA) (*ibid.*). This scenario would imply that AMHs were the producers of the Châtelperronian industry and occupied this region between ~44-40 kya cal BP, prior to the onset of the Protoaurignacian and possibly overlapping with late Neandertals producing Discoidal/Levallois technologies.

## 3. A synthetic review of Châtelperronian lithic technology

Châtelperronian lithic technology is arguably one of the most well-studied and well-described of any of Europe’s Middle to Upper Palaeolithic industries (Bricker and Laville, 1977; Boëda, 1991; Guilbaud, 1993; Pelegrin, 1995; Harrold, 2000; Connet, 2002, 2019; Bordes, 2003; Roussel and Soressi, 2006; Grigoletto et al., 2008; Bachellerie, 2011; Bordes and Teyssandier, 2011b; Roussel, 2011; Soressi, 2011; Aubry et al., 2012, 2014; Baillet et al., 2014; Roussel et al., 2016b; Bodu et al., 2017; Discamps et al., 2019; Rios-Garaizar et al., 2022). Here, we combine primary data and observations from two Châtelperronian assemblages (Les Cottés, US 06 and Quinçay, En) with descriptions in the published literature to provide a detailed review of this technology. The review is presented following a synthetic structure which combines individual criteria into seven well-established analytical divisions of lithic *chaînes opératoires*: percussor type and striking gesture [1] core initialisation and configuration [2], striking platform management [3], blank production [4], core maintenance procedures [5], core discard [6], and targeted blanks and tooling [7].

### 3.1 Percussor type and striking gesture

Percussion techniques within the Châtelperronian can be generally summarised as reflecting the variable use of marginal – and, to a lesser extent - internal percussion, utilising a mineral (i.e. stone) hammer. The generally small thickness of blade platforms (~2-5mm), frequent soft abrasion of the external platform edge, ~90 exterior platform angles, common occurrence of splintered bulbs (*esquillement de bulbe*), weakly developed bulbs of percussion, and low frequency of pronounced internal platform ‘lips’ are consistent of a marginal application of force using a soft-stone hammer for *plein debitage* (e.g. Pelegrin, 1995; Bachellerie, 2011; Roussel et al., 2016; Bodu et al., 2017; Rios-Garaizar et al., 2022; Rios-Garaizar et al., 2012; Michel et al., 2019; Aubry et al., 2012; Aubry et al., 2014). Internal hard-hammer percussion is reported however, specifically for larger initialisation/maintenance products (Pelegrin, 2011) or the discrete production of blanks for end-scrapers (Michel et al., 2019). With this said, some experimental work has criticised the unequivocal distinction between hard or soft stone hammers (Roussel et al., 2009). Taking this into account, a more cautious interpretation would be that Châtelperronian knappers generally utilised a marginal striking gesture and a percussor of mineral nature for primary debitage (*plein debitage*) phases.

### 3.2 Core initialisation and configuration

Large blocks or slabs, and also large flakes, appear to be preferentially selected for Châtelperronian blade production. Given the character of discarded cores, it is likely that morphologies which afford the unproblematic installation of a wide and flat flaking surface are particularly desirable. Initialized Châtelperronian blade cores often show an installation of a one-sided crest (**Figure 1**), generally on a narrow face, prepared with unidirectional transverse removals which extend onto an adjacent wide surface (Bachellerie, 2011; Roussel et al., 2016b; Bodu et al., 2017). Two-sided starting crests also occur, but appear to be less frequent (**Figure 1; b**) (e.g., Connet, 2019). The second wide and narrow faces often show minimal preparation, although postero-lateral cresting does feature on some discarded blade cores. This shaping procedure often produces initialized cores with an asymmetric cross-section and the first generation of crested blades also often display this asymmetry (i.e. show laterally-steeped cross sections). Bladelet cores, which are produced on small blocks or large flakes (*debitage sur tranche)*, can show very similar initialisation procedures (asymmetric configuration of narrow and wide surface) – and crested lamellar elements (<13mm) occur in some Châtelperronian assemblages (Roussel and Soressi, 2013; Bachellerie, 2011; Roussel, 2011; Rios-Garaizar et al., 2022) (**Figure 1; a**). It must be noted however that there is variability in the character of bladelet core initialisation and configuration between assemblages – with some showing minimal preparation procedures (Bodu et al., 2017).

**Fig. 1.**
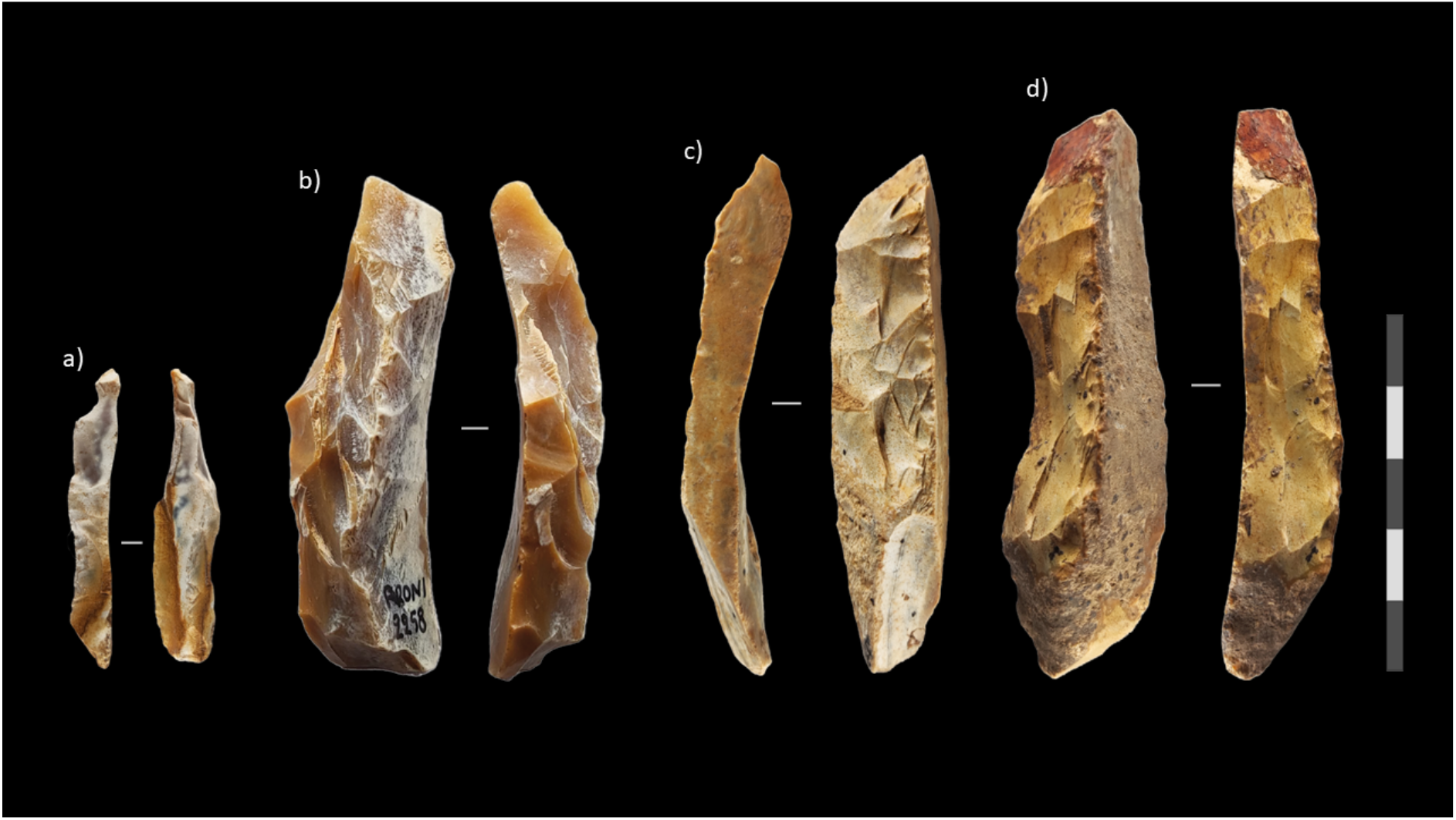
Châtelperronian crested blades (b-d) and bladelet (a) from initialisation/configuration stages. One-sided crests (a, c-d) and two sided-crest (b). Note the lateral-steeped cross sec7on (c). Artefacts are from the Châtelperronian of Quin*ç*ay (c, d) and Les CoE*é*s (a,b).

### 3.3 Striking platform management

The placement of a striking platform through the detachment of a large flake in the axis of the first debitage surface makes it possible to control the desired angulation, which is revived through the detachment of subsequent partial or total core tablet products. The angle between the platform(s) and the flaking surface(s) is nearly always between 75 and 90 degrees (Bachellerie, 2011; Roussel et al., 2016b; Bodu et al., 2017; Rios-Garaizar et al., 2022) (**Figure 2; a,b**). Two opposing platforms appear to be relatively frequently opened in early stages of core reduction/initialization (Bachellerie, 2011; Bodu et al., 2017). This is made evident by large, overshot blades which preserve an opposing (and often separated) platform on their distal end (**Figure 3; b-c**). However, exclusive unidirectionality is reported at some sites (e.g., Michel et al., 2019). Clear platform faceting is rare, if not entirely absent, from debitage – although some degree of discrete platform faceting is reported from some assemblages (e.g., Rios-Garaizar et al., 2012). More commonly, platforms are almost exclusively unmodified (**Figure 2; a-b**), excluding the relatively common occurrence of a soft abrasion on the external platform edge – likely related to the use of a soft-stone percussor. The maintenance of a near-perpendicular (75-90 degree) exterior platform angle, through successive tablet removals (total and partial), is a common feature however – with acute exterior platform angles on both cores and debitage being very uncommon (Bachellerie, 2011; Connet, 2019; Bodu et al., 2017; Michel et al., 2019; Porter et al., 2019).

**Fig. 2.**
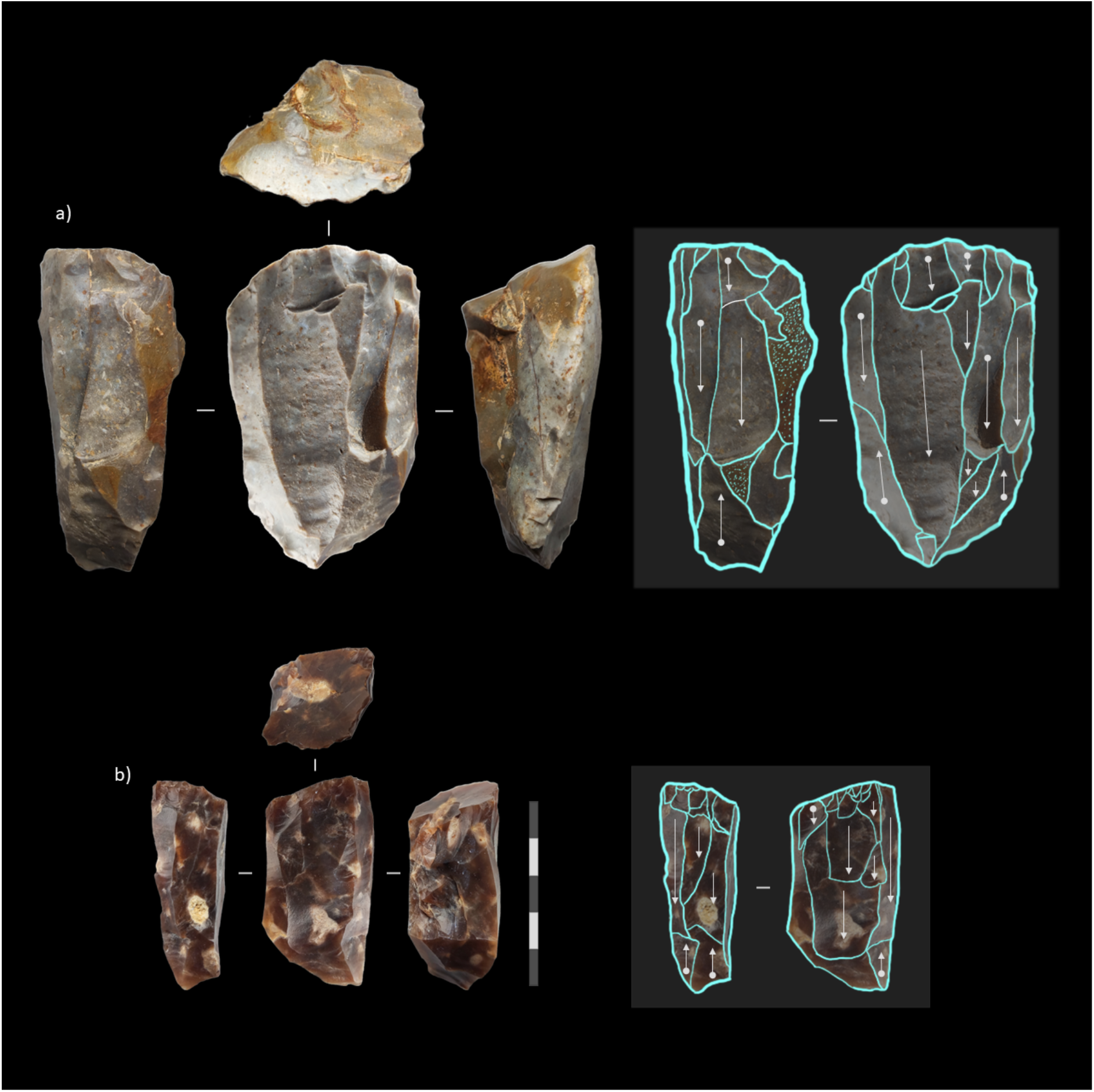
Châtelperronian blade (a) and bladelet core (b) showing postero-lateral cres7ng, plain plaForms, and EPA’s of ~75-80 degrees. Note the highly similar configuration and asymmetrical cross-sec7ons of the cores. Opposed and opposed and separated striking plaForm has been used for both maintenance and debitage procedures. Artefacts are from the Châtelperronian of Quin*ç*ay (a) and Les CoE*é*s (b).

**Fig. 3.**
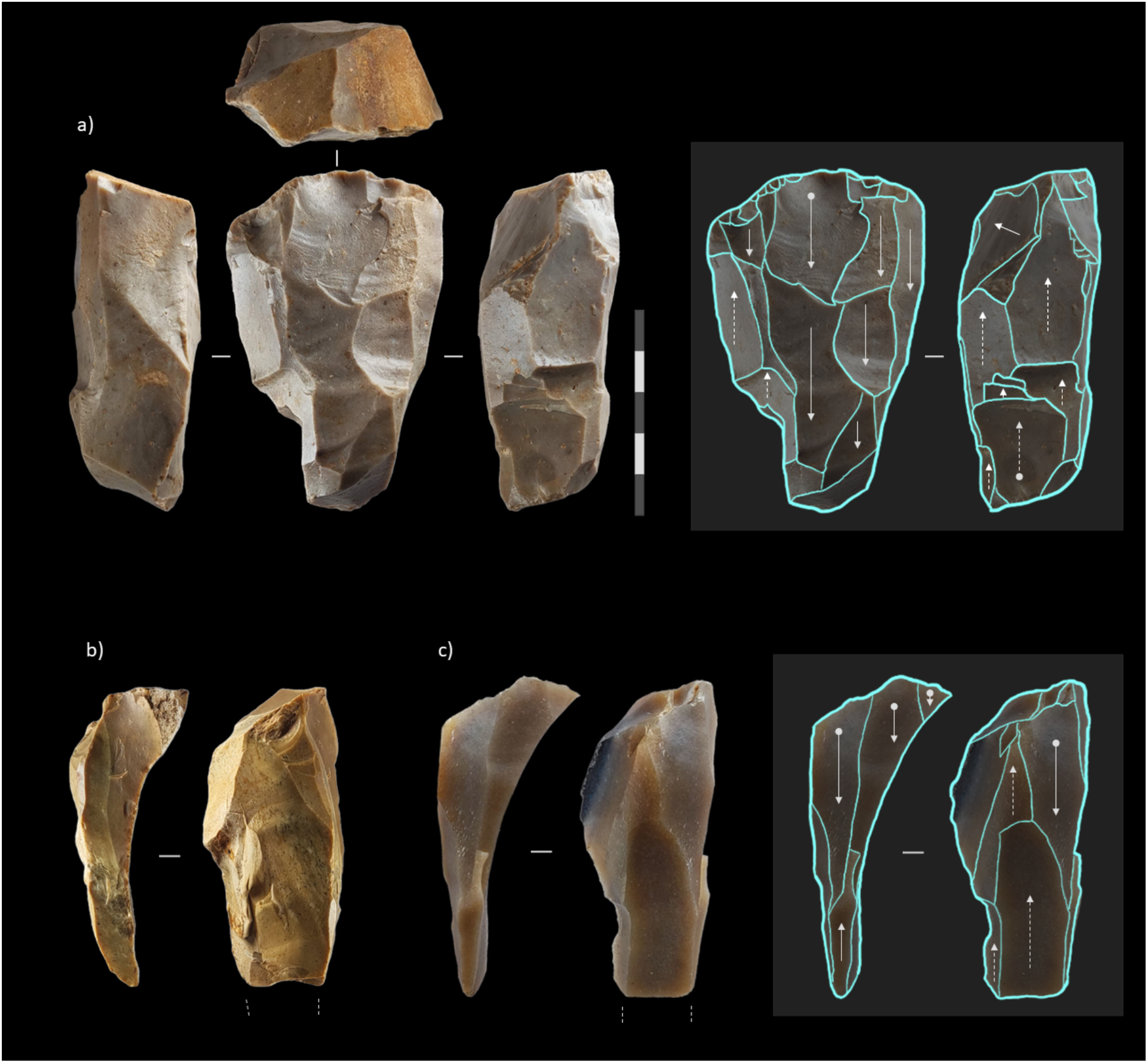
Discarded Châtelperronian blade core (a) and two overshot blades preserving an opposed and separated striking plaForm (b-c). Note the shiR to an opposed and separated striking plaForm for the final removals (dashed arrows) and the internal striking gesture applied to detach the last series of products (including the overshot blades themselves). Artefacts are from the Châtelperronian of Quin*ç*ay (a-b) and Les CoE*é*s (c).

### 3.4 Blank production

Large and medium sized blades are detached following a sub-parallel reduction method most often from the wide flaking surface of a core (e.g., Connet, 2019) (**Figure 3; a**). However, the narrow surface may sometimes play a secondary role, and cores may be entirely re-oriented at the very late stage of reduction (e.g., Roussel et al., 2016) (**Figure 3; a**). Some cores, as at Quinçay and Les Cottés, show a debitage progression on as many as three distinct (i.e. non-continuous) surfaces - often one wide and two narrow – producing a distinct rectangular cross section in core morphology (Roussel, 2011; Roussel et al., 2016b).

When blades are detached from two opposed striking platforms, blank production commonly follows short sets of unidirectional removals from a single platform before a switch to an opposed or opposed and separated platform. In this sense, when reduced using two opposed platforms, Châtelperronian blade reduction does not reflect true intersected bidirectionality, but rather a form of alternating unidirectionality (Roussel, 2011). Within the debitage, this procedure is witnessed in the co-existence of blanks with unidirectional and bidirectional dorsal scar patterns – the latter of which are often in lower proportions (Bachellerie, 2011; Roussel, 2011; Bodu et al., 2017; Rios-Garaizar et al., 2022) – as well as in overshot blades preserving an opposing striking platform (**Figure 3; b-c**). This is not ubiquitous however, with some assemblages strongly indicating the exclusive use of unidirectional reduction methods (Michel et al., 2019; Grigoletto et al., 2008). Blanks with lateral-steeped cross sections are often detached from either a) the intersection between two flaking surfaces or b) the intersection between the flaking surface and the core edge or back (during late-stage reduction). In some cases, and likely during the later stages of blank production, reduction is shifted to an opposed and separated striking platform prior to the discard of the core (e.g., Roussel et al., 2016) (**Figure 3; a**).

Small blade and bladelet production (**Figure 2; b**, **Figure 4; a**) appears to be quite variable, evidenced by the presence of at least three modalities leading to the production of small laminar elements: ‘simple’ burin-type cores on the edge of flakes and blades (Pelegrin, 1995; Roussel, 2011; Aubry et al., 2012, 2014; Bodu et al., 2017), prismatic/volumetric bladelet cores (Roussel, 2011; Roussel et al., 2016; Rios-Garaizar et al., 2022), and a continuum of reduction from blades to bladelets on the same core (Bachellerie, 2011; Floss et al., 2016). For burin-type cores, products are initially detached transversally from the distal edge of a thick flake, with production often progressing to the ventral face of the flake or blade (e.g., Pelegrin, 1995; Bodu et al., 2017). These cores generally however show a relatively low degree of productivity, and are often abandoned due to recurrent hinge fractures after a short series of removals. This may be related to the steep angle between the platform and flaking surface in combination with an insufficiently marginal striking gesture.

**Fig. 4.**
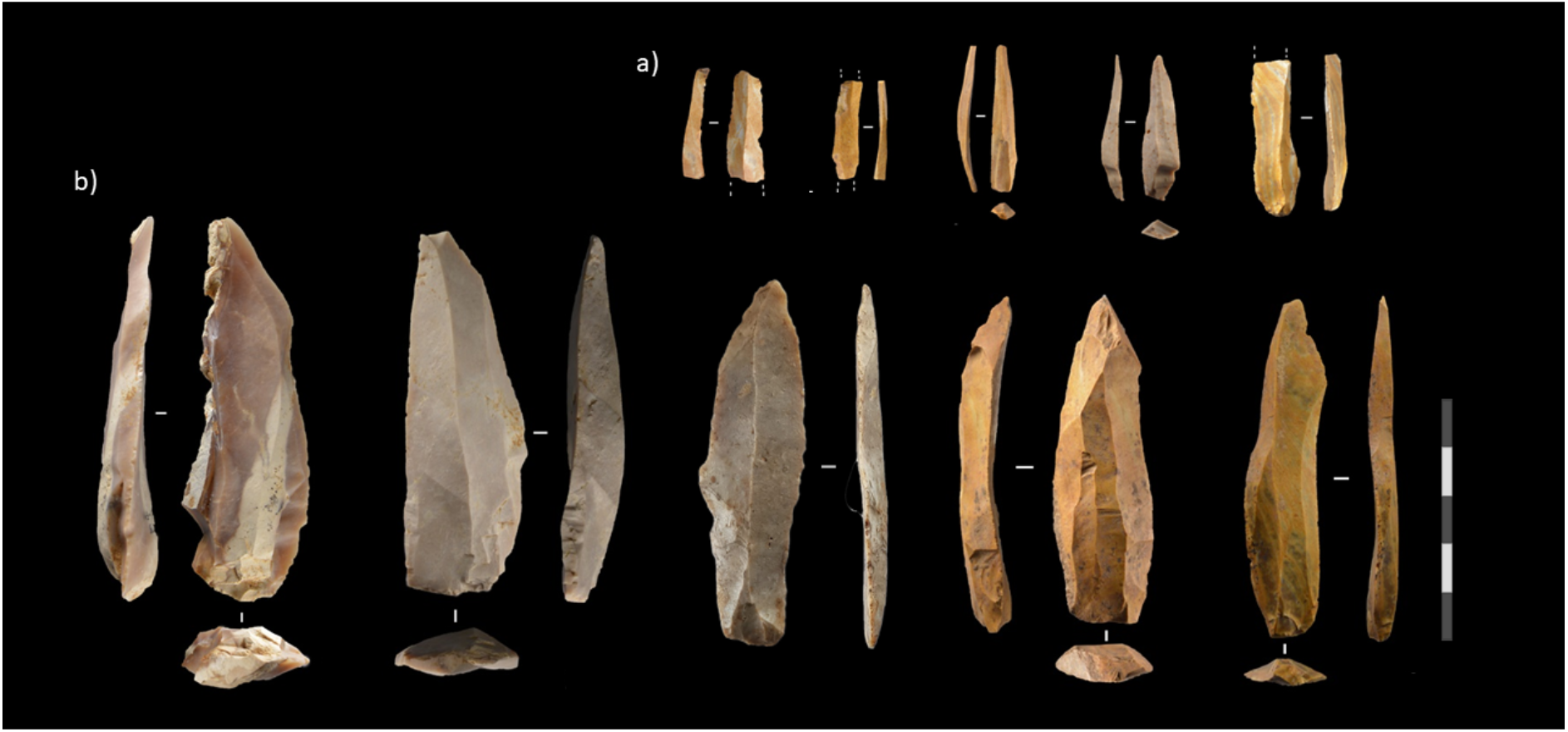
Bladelets (a) and blades (b) from *plein debitage* produc7on stages. Artefacts are from the Châtelperronian of Quin*ç*ay and Les CoE*é*s.

Volumetric bladelet cores are generally more productive and mirror, to a good extent, the method used for large-medium blade reduction (Pelegrin, 1995; Bachellerie, 2011; Roussel et al., 2016b; Bodu et al., 2017). Production similarly appears to be aimed at the obtention of straight or slightly curved parallel-edge bladelets (**Figure 4; a**). This appears to be the case in almost all Châtelperronian assemblages where bladelet production has been reported. One notable and clear exception to this is a highly productive, convergent bladelet core from the recently published site at Aranbaltza, Spain which – for the moment – remains largely unique within well-described Châtelperronian assemblages (Rios-Garaizar et al., 2022).

### 3.5 Core maintenance procedures

Blade production is maintained through the use of second-generation crested blades, debordant blades, core tablet removals, and laminar rejuvenation flakes. Neo-crested blades are a common maintenance procedure applied throughout core reduction to restore lateral convexities (**Figure 5; b-e**). Similar to initial crests, these are generally installed at the intersection of two flaking surfaces – producing a crested blade with an acute triangular cross section. Neo-crested blades can show substantial size variability, indicating that this may be a recurrent maintenance procedure throughout the reduction of a core. The removal of non-crested debordant blades with a laterally-steeped cross section likely fulfilled a similar role when cresting was not required to alter the angle and morphology of the removal (**Figure 5; f**).

**Fig. 5.**
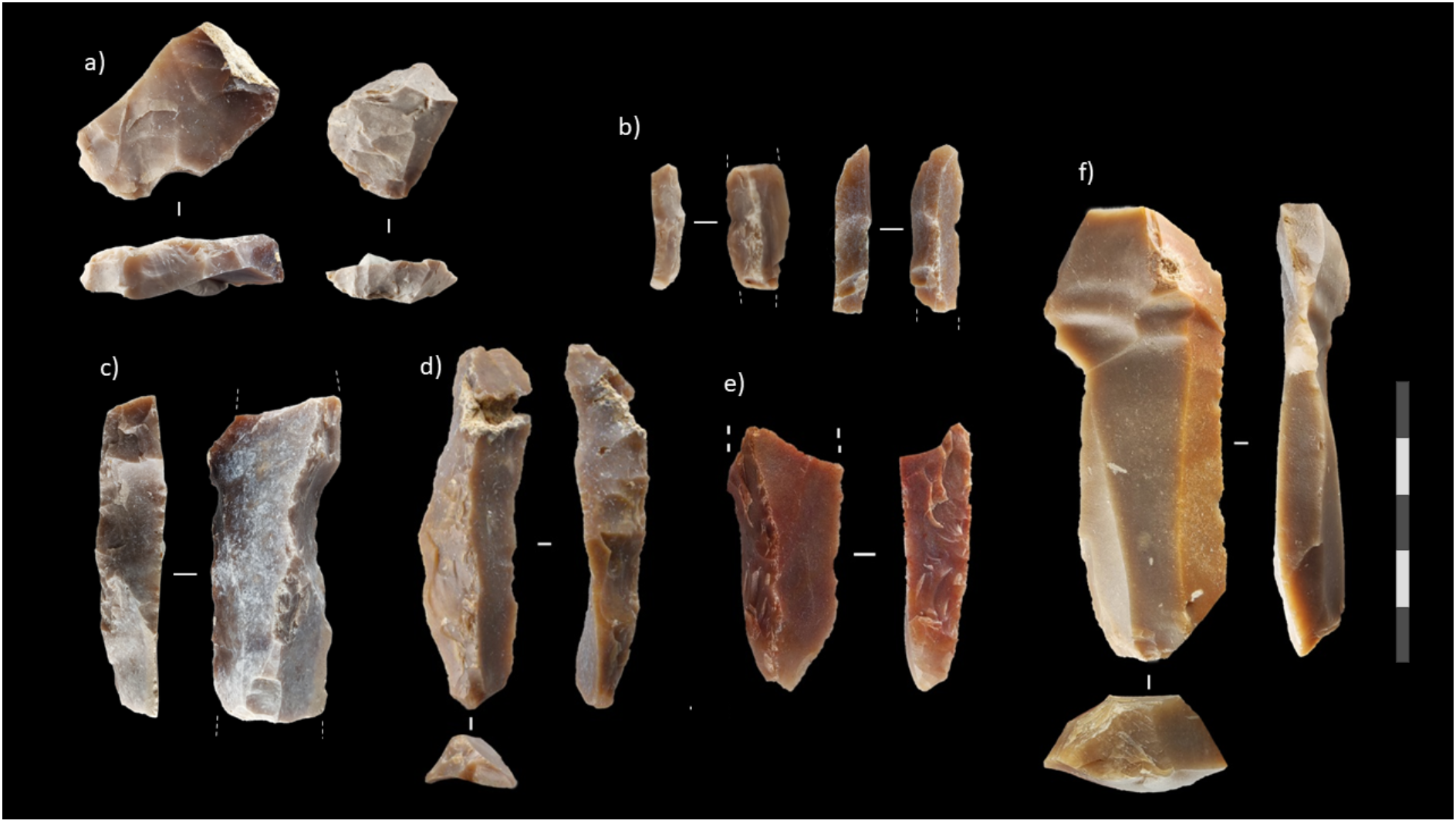
Products rela7ng to maintenance interven7ons. a) Core tablet removals, b-e) neo-crested products of various sizes, f) large debordant blade. Artefacts are from the Châtelperronian of Quin*ç*ay (d) and Les CoE*é*s (a-c, e-f).

The presence of partial or complete tablet removals on blade cores shows that rejuvenation of the striking platform was a relatively common maintenance procedure (**Figure 5; a**), an observation which is supported by the identification of core tablets corresponding to blade production in almost all Châtelperronian assemblages. For example, at Les Cottés, maintenance tablets are numerous – and are either detached frontally or, less often, laterally to the main flaking surface. The counter-bulbs present on the lateral edges of the tablets indicate debitage sequences of wide blades, narrow blades, and bladelets.

Laminar rejuvenation flakes are large flakes removed from the wide flaking surface of blade cores, generally with an internal striking gesture, after the surface has become flattened due to the extraction of several blades (e.g. Roussel et al., 2016; Bodu, 1990) (**Figure 6; c**). This procedure removes nearly the entire flaking surface, serving to construct new transversal convexities. Following the detachment of this flake, an asymmetrical blade is removed from the intersection of the narrow and wide surface and blade production is recommenced. These products are often converted into end-scrapers, as is evident by the presence of numerous laminar negatives on the dorsal face of many Châtelperronian end-scrapers (Baillet et al., 2014; Roussel et al., 2016b) (**Figure 6; c**).

**Fig. 6.**
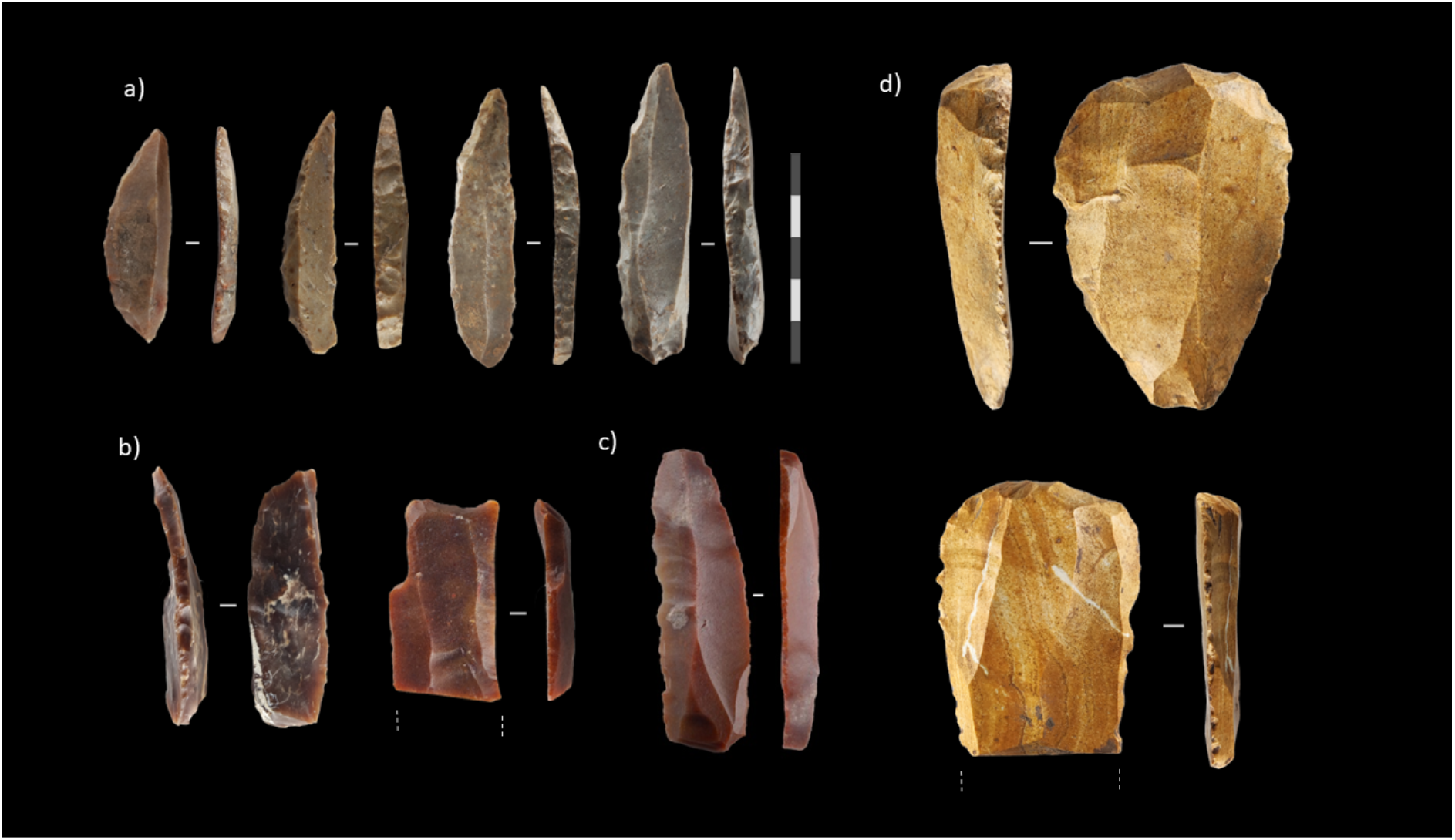
Examples of retouched Châtelperronian tools. a) Curved-backed points/knives (Châtelperronian points), b) burins (dihedral and on trunca7on), c) wide-fronted end-scrapers produced on large laminar rejuvenation flakes. Artefacts are from the Châtelperronian of Quin*ç*ay (a,c) and Les CoEés (b).

### 3.6 Core discard

A recurring and notable feature of discarded blade and bladelet cores in the Châtelperronian – which can in some sense in fact be considered quite typical of the industry – is a series of deep hinged removals affecting one or more of the flaking surfaces (Roussel et al., 2016; Bachellerie and Bordes, 2018). This may relate to the utilisation of a more internal striking gesture in the final phases of core exploitation – as may be evident in the pronounced *contra bulbs* visible in the negatives of final removals. Some of which has been proposed as indication of apprenticeship in flint-knapping (Bachellerie and Bordes, 2018).

### 3.7 Targeted blanks and tooling

Châtelperronian blade core reduction is targeted near-exclusively at the obtention of one product: regular blades for the manufacture of the arch-backed points which typify this industry (**Figure 6; a**). The targeted blanks are straight, sub-parallel blades predominantly between 35-85mm in length, 12-35mm in width and 4-9mm in thickness. Additionally, naturally-backed blades detached are sought after for the production of Châtelperronian points.

Second-choice blades are frequently modified by a lateral retouch (marginal or abrupt), or converted into simple dihedral burins (**Figure 6; b**), while more robust blades and technical flakes detached from the wide flaking surface of cores are converted to end-scrapers (e.g., laminar rejuvenation flakes) (**Figure 6; c**). At some sites, the discrete production of elongated flakes has been identified – linked to the manufacture of large-fronted end-scrapers (Michel et al., 2019). The Châtelperronian toolkit consists near-exclusively of ‘Upper Palaeolithic’ tool forms produced on products deriving from well-developed laminar core reduction methods (**Figure 6**). Lamellar reduction seeks the production of relatively regular, sub-parallel bladelets – which are most often left unretouched. In some contexts, these unretouched lamellar products show evidence of use (Bodu et al., 2017). When retouch is present, oblique truncations – somewhat mirroring the retouch of Châtelperronian points – are present. In addition, direct or inverse lateral retouch on bladelets is observed at several sites – including Les Cottés, Quinçay, and Aranbaltza II (Soressi et al., 2006; Bachellerie, 2011; Roussel, 2011; Talamo et al., 2012, 2020; Roussel et al., 2016; Bodu et al., 2017; Discamps et al., 2019; Rios-Garaizar et al., 2022).

### 3.8 Summary

Châtelperronian lithic technology reflects a well-developed laminar reduction system which is initiated and maintained through systematic cresting procedures (antero-lateral, postero-lateral). Two platforms are commonly opened, and blades are detached in alternating series. The idealised core reduction method can be considered as a variant of ‘asymmetric core reduction’ (Roussel, 2011; Roussel et al., 2016; Zwyns, 2021). *Plein debitage* is most likely detached utilising a soft-stone percussor and more marginal percussion, while secondary products (technical flakes, debordant blades etc.) can show signs of a more internal percussion gesture and are often converted into end-scrapers or cores for small lamellar elements (bladelets). Arch-backed points (Châtelperronian points) are often the most numerous, and most distinctive, retouched element within Châtelperronian assemblages. Their size can vary widely, from near-bladelet sized examples to robust and heavily-backed examples. The rest of the retouched toolkit consists near-exclusively of ‘Upper Palaeolithic’ tool forms (end-scrapers, ‘simple’ dihedral burins, laterally-retouched blades etc.). The production of bladelets in this industry, including rare retouched bladelets, has now been demonstrated at multiple sites and in multiple modalities.

## 4. Prospects and pitfalls of an intrusive model for the Châtelperronian stone tool industry

This review has reinforced that the Châtelperronian is a distinctive, fully ‘Upper Palaeolithic’ (i.e. laminar), and well-understood lithic industry. Questions concerning the nature, origins, and makers however have been the source of ongoing discussion and debate for many years (e.g. D’Errico et al., 1998; Bar-Yosef and Bordes, 2010; Caron et al., 2011; Ruebens et al., 2015; Roussel et al., 2016b; Hublin et al., 2020; Djakovic et al., 2022; Rios-Garaizar et al., 2022). Here, we present a point-by-point evaluation of some prospects and problems for an intrusive model for the Châtelperronian and the onset of the Upper Palaeolithic in France and northern Spain.

### 4.1 Some prospects for an intrusive model of the Châtelperronian

#### A Northern Early Ahmarian origin of the Châtelperronian resolves the issue of ambiguous local continuity

The local origin model of the Châtelperronian has been a source of debate for many years. At present, the biggest empirical issues pressuring this model are stratigraphic, technological, and chronological. It is increasingly appearing that the MTA Type-B is separated from the Châtelperronian by at least a Discoidal, and likely also a Levallois, phase at sites where they occur together (Jaubert, 2011; Jaubert et al., 2011; Gravina and Discamps, 2015; Gravina et al., 2022). This is not consistent with a continuity of the Châtelperronian from this Mousterian facies. Furthermore, the structure of Châtelperronian lithic technology is totally inconsistent with the preceding (flake-based) Levallois and Discoidal phases (see Faivre et al., 2017). A direct external origin of this industry would in effect resolve the issue of the sudden appearance and spread of a well-developed, volumetric, crest-based technology (as described above) – releasing the Châtelperronian from increasingly ambiguous arguments pertaining to a local techno-cultural precursor.

There are a number of ways to interpret technological changes in the archaeological record of the Middle to Upper Palaeolithic transition beyond the two most common approaches (local continuity and geographically continuous migration) (**Figure 7; a,b**). For example, a migration can be geographically discontinuous – i.e. ‘Invisible’ between the nodes – due to maritime voyages (**Figure 7; d**). Alternatively, a small and highly mobile migrating population is unlikely to leave a dense and recognizable lithic footprint between the areas and/or sites which they occupied more intensely and for a longer duration. For example, we can consider the recently published Micocquian expression at Chagrskaya Cave in the Altai foothills – roughly 3000km away from the suggested geographic distribution of this industry (Kolobova et al., 2020). This appears to be an example of a long-distance population movement associated with a discrete lithic tool-kit – which is in fact supported by genetic evidence (*ibid.*). It is a good reminder that the migration of highly mobile hunter-gatherer groups can remain archaeologically invisible between the central nodes – instead perhaps reflected by assemblages exhibiting mixed characteristics and typo-technological features labelled as one industry or another. A large geographic distance between similar archaeological assemblages does not, and should not, necessarily preclude a direct connection/relationship between those assemblages. Conversely, a similar geographic distribution between two sets of assemblages is not necessarily evidence for a connection/relationship between these assemblages – for example, the MTA-B and Châtelperronian (Gravina and Discamps, 2015).

**Fig. 7.**
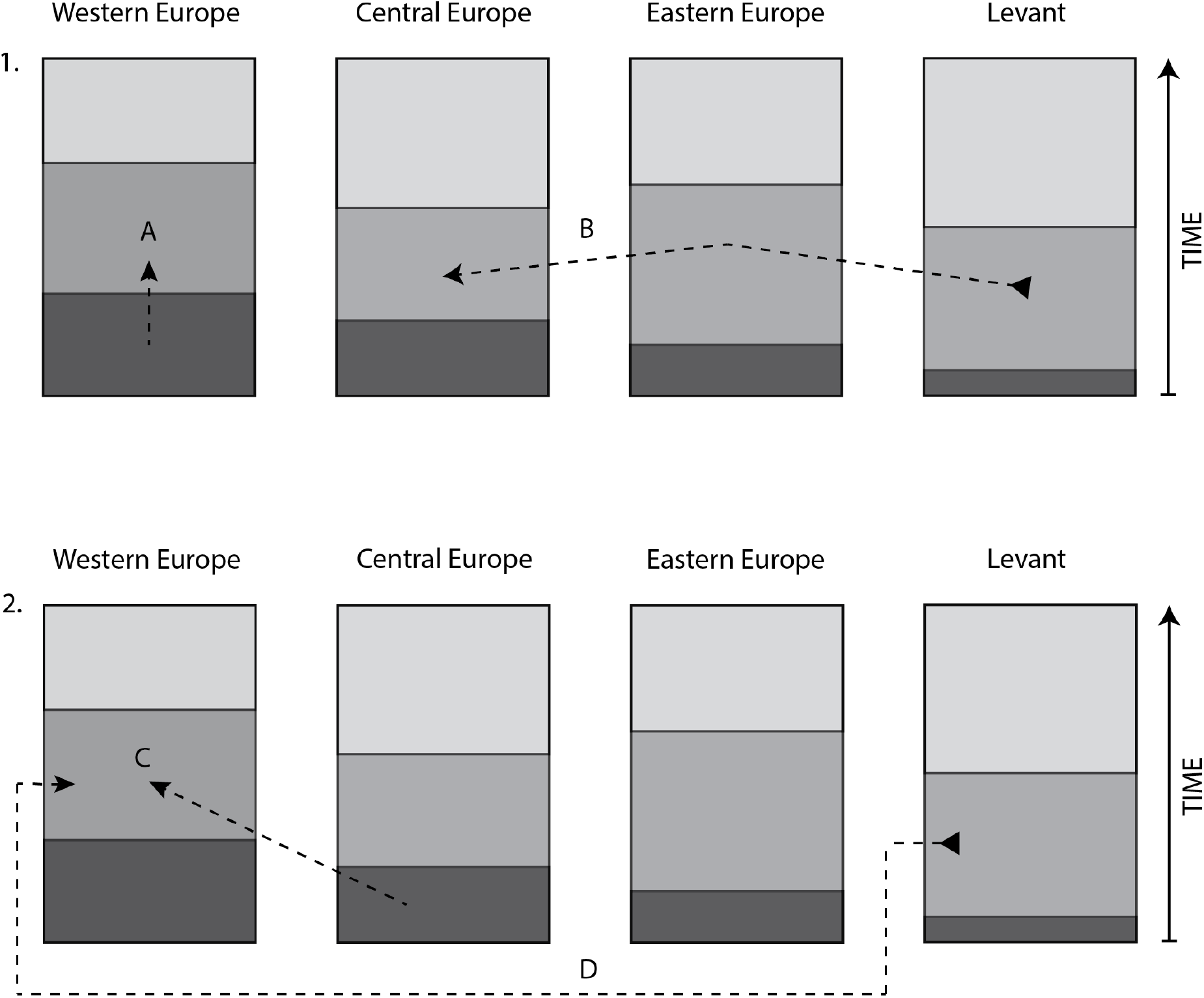
Simplified illustra4on of two common (1) and two uncommon (2) interpretative models for explaining the spread/emergence of archaeological industries relating to the Middle to Upper Palaeolithic transition. 1. A) origin via evolution from a local substrate (e.g. Châtelperronian from MTA-B), B) intrusive technology via geographically con4nuous migra4on (e.g. Protoaurignacian spread from Levant to western Europe). 2. C) origin from evolution from non-local substrate, D) intrusive technology via geographically discontinuous (or archaeologically ‘invisible’) migra4on.

Concerning the elongated production characterizing the MTA Type-B, one may wonder whether this industry may actually instead have some form of relevance to the Neronian (IUP) industry, as it may in fact share a similar chronological position - both industries likely occupying a period of time between ~56 and 50 kya cal BP (Soressi et al., 2013; Gravina and Discamps, 2015; Slimak et al., 2022).

#### The earliest available dates for any Protoaurignacian or Initial Upper Palaeolithic context in Europe currently come from southern France and northern Spain

A prediction of the ‘Three Waves’ model for the onset of the Upper Palaeolithic in France and northern Spain is that the related industries should occur earlier in the Mediterranean region than in other regions of Europe (Slimak, 2023). For the IUP and Protoaurignacian, this is in fact consistent with currently available data. The IUP (Neronian) at Grotte Mandrin and the Protoaurignacian at sites such as Isturitz and Gatzarria are currently the earliest dated occurrences of these respective industries in any region of Europe (Barshay-Szmidt et al., 2012, 2018, 2020; Djakovic et al., 2022; Slimak Ludovic et al., 2022). This pattern, while acknowledging sample size and biases in research history, is consistent with the pattern which is predicted under the ‘Three Waves’ model or similar models. For the Châtelperronian, this industry often appears to share a geological correlation/association with a Discoid or Levallois Mousterian when they are found in the same stratigraphic context (e.g. Aubry et al., 2012, 2014; Rocca et al., 2017; Gravina et al., 2018) – although it appears to post-date the IUP at both Grotte Mandrin and Bacho Kiro based on C14 measurements (Hublin et al., 2020; Slimak Ludovic et al., 2022).

#### If the Châtelperronian has a direct origin in the Levant, it is possible that small quantities of Levantine raw material (e.g. flint) could be found in Châtelperronian assemblages

Arguably the most direct way to demonstrate a Mediterranean connection between the Levant and France during the time of the Châtelperronian/Northern Early Ahmarian is to identify raw material (e.g. flint) from the Levant in a Châtelperronian assemblage (or used to produce a Châtelperronian point). This would directly demonstrate the physical transport of material from the Levant to western Europe, although it would not necessarily imply the crossing of the Mediterranean Sea (nor would it demonstrate a direct connection between the two aforementioned industries). Such a demonstration would in any case represent a ‘needle in a haystack’-type of discovery. With this said, despite the limitations, raw materials of unknown origin within Châtelperronian (and Protoaurignacian?) assemblages in France may prove to be key to demonstrating the viability of different explanatory models for AMH migration into western Europe in the future.

#### Some technological similarities between the Châtelperronian and Protoaurignacian are expected under a ‘common origin’ model, and appear to be observed in reality

Of relevance to the ‘Three Waves’ model is the acknowledged and increasing presence of similar bladelet technologies, osseus artefacts, and personal ornaments within a number of Châtelperronian and Protoaurignacian contexts (e.g. Julien et al., 2019; Roussel et al., 2016; Rios-Garaizar et al., 2022) – which would be expected under a ‘shared cultural substrate’ model such as ‘Three Waves’. Regardless, it has been established that the Châtelperronian is technologically more similar to the Protoaurignacian than it is to any local late Mousterian industries (Bordes and Teyssandier, 2011; Gravina and Discamps, 2015; Roussel et al., 2016). The principal technological differences between these industries are those of core external platform angle, unidirectionality/bidirectionality, knapping gesture, and primary debitage intention (laminar in the Châtelperronian versus lamellar in the Protoaurignacian) (e.g. Roussel et al., 2016). On the other hand, the technological similarities between these industries include: cresting technology, unmodified platforms, fully laminar structures, soft-stone percussion, and ‘Upper Palaeolithic’ tool forms (e.g. end-scrapers, retouched blades, burins etc.) (Djakovic et al., 2022). Curiously, it appears that the primary differences between debitage trends identified for the Northern Early Ahmarian and Southern Early Ahmarian mirror to a notable extent those observed between the Protoaurignacian and Châtelperronian (Kadowaki et al., 2015). Relevant to this discussion is the observation that some Châtelperronian assemblages – such as Aranbaltza II - show a pronounced and well-developed bladelet component (Rios-Garaizar et al., 2022) featuring in our opinion potential technological and typological (Dufour bladelet) affinities with the regional Protoaurignacian.

#### An industry with the same set of technological features as those that characterize the Châtelperronian does not appear to have been produced, in the current state of knowledge, by any other Neandertal groups

This statement has nothing to do with discussions of ‘cognitive capacity’ or ‘behavioral modernity’, but simply serves to highlight the fact that the Châtelperronian appears to be a technological and behavioral outlier when compared to all other industries presently associated with/attributed to Neandertals. Of course, this pattern is largely a construction of research history - and could very well be rendered incorrect or irrelevant with new discoveries and/or re-evaluations. Nonetheless, the point remains that there presently exists no behavioral analogue to the Châtelperronian in the broader Neandertal techno-cultural repertoire. Conversely, the Châtelperronian does share typo-technological similarities to certain lithic complexes of the MSA in Africa (for example, the Howiesoon’s Poort complex) as well as the Northern Early Ahmarian of the Levant (Soriano et al., 2007; Kadowaki et al., 2015; Slimak, 2023).

### 4.2 Some problems for an intrusive model of the Châtelperronian

#### There are numerous Neandertal fossil remains in association with Châtelperronian artefacts at Grotte du Renne, Arcy-sur-Cure (France)

The large suite of paleoanthropological evidence from the Châtelperronian levels at Grotte du Renne, Arcy-sur-cure includes exclusively Neandertal skeletal remains (Hublin et al., 1996; Bailey and Hublin, 2006). Irrespective of opinions concerning the integrity of the site, this evidence cannot simply be dismissed. Models of the nature, origins and authors of the Châtelperronian should be based on constructive discussions of empirical evidence - and it remains the case that not a single confirmed *Homo sapiens* fossil remain has yet been discovered in association with Châtelperronian artefacts. This must of course be balanced with the fact that human remains are in general very scarce within this period. However, this if anything makes the suite of Neandertal skeletal material from the Grotte du Renne all the more remarkable. Interestingly, a Neandertal tooth directly-dated to between 43-42 kya cal BP is also reported from the Protoaurignacian layer (US 04inf) at the site of Les Cottes (Hajdinjak et al., 2018), but a detailed analysis of its context is yet to be published.

#### The chronological support for a Northern Early Ahmarian origin of the Châtelperronian is ambiguous at best

For the Levantine origin model to be correct, it would be necessary that the Northern Early Ahmarian chronologically pre-dates – at least partially – the Châtelperronian of France and northern Spain. There is, in the current state of the art, no clear support for such a scenario (e.g. see Kadowaki et al., 2015; Djakovic et al., 2022). Calibrated radiocarbon ages from the Northern Early Ahmarian of Ksar Akil and Üçağɩzli provide ranges between ~41 and 34 kya cal BP (Kuhn et al., 2009; Douka et al., 2013), placing them either briefly contemporaneous with or post-dating the accepted temporal range of the Châtelperronian (i.e. somewhere between 44-40 kya cal BP). With this said, the Northern Early Ahmarian of Kebara has provided calibrated radiocarbon ages ranging between ~50 and 39 kya cal BP (Rebollo et al., 2011) – which could represent an example of an assemblage associated with this industry pre-dating the emergence of the Châtelperronian around 44 kya cap BP. It should be noted, however, that the production and comparison of robust chronologies for these industries is hindered by the fact that this period is situated near the upper acceptable age limit for radiocarbon dating. This is perhaps further exacerbated by the construction of chronologies based on the dating of different sample types (bone, charcoal, shell etc.).

#### The similarity/dissimilarity of Châtelperronian and Northern Early Ahmarian lithic technology has not been sufficiently demonstrated

One of the most obvious requirements for the proposition of a Northern Early Ahmarian origin model for the Châtelperronian is a detailed demonstration that these lithic industries and their associated technological behaviors are effectively analogous. This has not as of yet been demonstrated, although morpho-typological similarities between the retouched backed points of the Northern Early Ahmarian and the Châtelperronian are evident based on published illustrations and were in fact already stressed in the previous century (see discussions in Kadowaki et al., 2015; Slimak, 2023). This is made further interesting when considering that the mean size (length and width) of backed points from Ksar Akil (Layer XVII, n=75) (Bergman, 1981) and Kebara (Layer E, n=78) (Ziffer, 1978) form a tight cluster with that of the Châtelperronian points recovered from Quinçay (Layer En, n=77) (Roussel et al., 2016) (**Figure 8**) when compared to those of the Southern Early Ahmarian complex (Kadowaki et al., 2015).

**Fig. 8.**
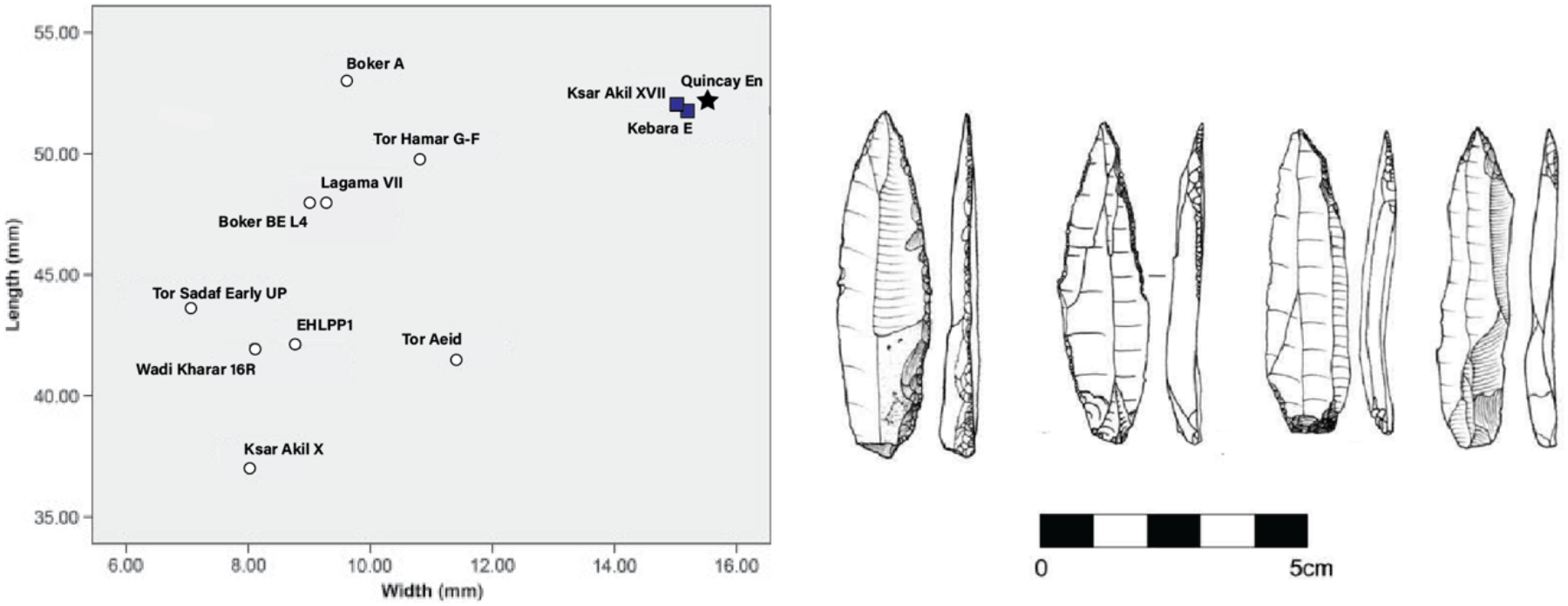
LeR: ScaTerplot showing mean length and width of points from the Southern Early Ahmarian (white circles), Northern Early Ahmarian (dark squares), and Châtelperronian of Quinçay (black star) (modified aRer Kadowaki et al., 2015). Kebara E (n=78), Ksar Akil XVII (n=78), Quincay En (n=77). Right: Example of Ahmarian retouched points from Üçağɩzli cave (modified aRer Kuhn et al., 2009).

To test the viability of a direct relationship between these industries, it would be at a minimum necessary to demonstrate that the Châtelperronian is substantially more similar (technologically) to the Northern Early Ahmarian than to other contemporaneous ‘initial’ Upper Paleolithic industry in Europe. For example, the Lincombian-Ranisian-Jerzmanowician (LRJ) industry, which also represents a break with the preceding Middle Palaeolithic and - similarly to the Châtelperronian – is characterized by the production of blades from bidirectional cores utilizing marginal, and likely soft-stone percussion (Flas, 2011; Demidenko and Škrdla, 2023). Bladelet production has also been documented at some LRJ sites (Demidenko and Škrdla, 2023), and some cores (both blade and bladelet) show intriguing structural similarities to Châtelperronian counterparts (see for example Figure 5 in Flas, 2011). In this comparison, however, typological differences are immediately evident – with the LRJ typified by the production of large points characterized by flat, invasive retouch on one or both surfaces (Flas, 2011; Wiśniewski et al., 2022; Demidenko and Škrdla, 2023). It is important to highlight that numerous distinctive/diagnostic technological and typological categories exist within the Châtelperronian beyond the eponymous arch-backed point (for example, technical products, percussion techniques, and distinctive end-scrapers), which make a direct evaluation of typo-technological similarity between this industry and any potential analogues well within the realm of viability.

#### We have no physical evidence of sea-faring during the MP-UP transition, which is a behavior required by a direct Levantine origin model

There exists in this period no physical evidence of sea-faring craft or related technology of any kind, and in any region. However, there is ‘indirect’ evidence of sea-faring based on the arrival of *Homo sapiens* to Australia ~65 kya cal BP (Clarkson et al., 2017). Further complicating matters is the fact the sea level rises have likely inundated many archaeological sites which are relevant to these discussions. However, given the peopling of Australia, it is certainly not unreasonable to presume that some human groups had developed a decent level of seafaring capability by the MP-UP transition (~50-40 kya cal BP) – if not in fact considerably earlier. Modelling has shown that the accidental arrival of humans to Australia from the islands of Timor and Roti by drifting alone is very unlikely to have occurred (Bird et al., 2018), while genetic evidence indicates that Australia was colonized in a single phase – with very limited geneflow following this colonization – suggesting a founding population large enough to sustain long-term survival (Tobler et al., 2017). The colonization of Australia was most likely the result of a purposeful and coordinated series of maritime crossings – with the crossing to Australia being two or three times longer than the multiple previous crossings required to reach the islands of Timor and Roti (Bird et al., 2018; Norman et al., 2018). Direct evidence of such crossings (i.e. human-crafted maritime technologies) may however completely evade us for the foreseeable future due to a lack of preservation. Given the fragmented character of the archaeological record, we should remember that the absence of evidence is not evidence of absence. The colonization of Australia over 60 thousand years ago forces us to rethink both the importance and antiquity of maritime voyages in the spread of *Homo sapiens* across the globe.

## 5. On testing hypotheses for the onset of the Upper Palaeolithic in France and northern Spain

### sedDNA, ZooMs, and recovering DNA from bone tools and personal ornaments

The recovery of sedimentary DNA from bulk sediment and indurated blocks (Massilani et al., 2022), in combination with ZooMs analysis of non-identifiable faunal fragments from Châtelperronian contexts, will undoubtedly progress discussions concerning the makers and origins of this industry in the years to come. Additionally, the recent publication of a method for the non-destructive extraction of ancient human DNA directly from Palaeolithic ornaments and tools made of bone or tooth opens a new door towards connecting discrete hominin individuals with discrete archaeological artefacts (Essel et al., 2023). Given the emerging picture of demographic complexity in this period (e.g. Hublin et al., 2020; Hajdinjak et al., 2021; Prüfer et al., 2021; Vallini et al., 2022; Slimak Ludovic et al., 2022), it is likely also necessary to at least entertain the hypothesis of mixed groups (Compton et al., 2021; Stringer and Crété, 2022). However, it is critical that taphonomic approaches be implemented to assess the integrity of archaeological contexts (Texier, 2000; Bordes, 2003; Gravina et al., 2018) prior to the construction of higher level models and sedDNA analysis.

### Technological comparison of Châtelperronian, Northern Early Ahmarian, and contemporaneous European EUP technologies

To demonstrate the potential viability of a Levantine origin for the Châtelperronian, it would be necessary – at a minimum - to show that Châtelperronian lithic technology is significantly more similar to the Northern Early Ahmarian than it is to all contemporaneous European Early Upper Palaeolithic industries. A combination of *chaine opératoire,* attribute analysis, and 3DGM approaches would likely provide a highly informative evaluation of similarities and differences in technological behaviours at varying scales of observation (Zwyns, 2012, 2021; Falcucci et al., 2017; Kolobova et al., 2020; Archer et al., 2021). Concepts such as *near ubiquity* and *intricate complexity,* introduced to archaeological studies from primatological work (Byrne, 2007; Zwyns, 2021), could be used to make predictions about what kinds of technological behaviours would be necessary to effectively demonstrate a direct transmission of information between individuals within and between discrete archaeological assemblages. If the Châtelperronian is in fact explicitly analogous to the Northern Early Ahmarian, or any other industry, it should be demonstratable to a reasonably high level through an integrated technological approach.

### Raw material movements and geographic patterns

For an intrusive Levantine model, it would be useful to be able to demonstrate that the Châtelperronian sites nearer the Mediterranean coast are chronologically older than those further away from it. This is potentially the case for the Protoaurignacian (Barshay-Szmidt et al., 2020; Djakovic et al., 2022), but remains unclear for the Châtelperronian for various reasons. Raw material transport patterns within the Châtelperronian and Protoaurignacian could potentially help to infer patterns of population movement across the region. A key assumption of the Mediterranean origin hypothesis of these industries would be that the spread of these industries within western Europe was somewhat directional (e.g. moving from south to north). Some studies have already suggested that at a number of Châtelperronian and Protoaurignacian sites there appears to be only south to north patterns of movement when considering non-local raw materials (Discamps et al., 2014). This is currently an anecdotal but interesting pattern, and it may indicate that a wide-scale analyses of raw material transport would be a fruitful research area with respect to inferring potential movement patterns. It is likely that the dynamics underpinning the Middle to Upper Palaeolithic transition in the region cannot be understood without a consideration of the geographic structuring underlying the chronological and technological data.

### Returning to key sites and sequences

The re-evaluation of key sites and sequences will continue to be an important path towards a better understanding of the Middle to Upper Palaeolithic transition in this key region. One need only look at the results of recent work at a number of eponymous MP-UP sites across the region to understand the value in critical re-evaluations of key sequences (Bordes, 2003; Jaubert et al., 2011; Gravina and Discamps, 2015; Faivre et al., 2017; Gravina et al., 2018, 2018). In this vein, a suite of multidisciplinary work is currently in progress at the site of Quinçay (France) – a site with a pronounced bladelet component (Roussel et al., 2016) and one of two Châtelperronian sites from which personal ornaments have been published (Soressi and Roussel, 2014; Roussel et al., 2016; Welker et al., 2017). Interpretations of the Middle to Upper Palaeolithic transition based on sites excavated many decades ago should be - in general - treated with due caution and, in the absence of modern re-evaluations, more attention should be paid to recently excavated sites. This is particularly the case when constructing reliable interpretations pertaining to technological characteristics and, perhaps especially, fossil and/or sedDNA associations (Lévêque and Vandermeersch, 1981; Hublin et al., 1996, 2012; Gravina et al., 2018).

## Acknowledgements

I.D. thanks Leonardo Carmignani for many enthusiastic conversations, as well as Brad Gravina, Emmanuel Discamps and Marc Thomas for very fruitful and occasionally heated discussions during excavations at Le Moustier.

## Funding

This research is funded by the Dutch Research Council (NWO) ‘Neandertal Legacy’ grant (VI.C.191.070) awarded to M. Soressi.

## Author contributions

ID. and M.S. conceptualised the paper. I.D. wrote the manuscript and prepared the figures. I.D., M.R. and M.S. reviewed the manuscript and figures.

## Conflict of Interest

The authors declare no competing interests.

## Notes

### Competing Interest Statement

The authors have declared no competing interest.

## References

Archer, W., Djakovic, I., Brenet, M., Bourguignon, L., Presnyakova, D., Schlager, S., Soressi, M., McPherron, S.P., 2021. Quantifying differences in hominin flaking technologies with 3D shape analysis. Journal of Human Evolution. 150, 102912.

Aubry, T., Dimuccio, L.A., Almeida, M., Buylaert, J.P., Fontana, L., Higham, T., Liard, M., Murray, A.S., Neves, M.J., Peyrouse, J.B., Walter, B., 2012. Stratigraphic and technological evidence from the middle palaeolithic-Châtelperronian-Aurignacian record at the Bordes-Fitte rockshelter (Roches d’Abilly site, Central France). Journal of Human Evolution. 62, 116–137.

Aubry, T., Dimuccio, L.A., Buylaert, J.P., Liard, M., Murray, A.S., Thomsen, K.J., Walter, B., 2014. Middle-to-Upper Palaeolithic site formation processes at the Bordes-Fitte rockshelter (Central France). Journal of Archaeological Science. 52, 436–457.

Bachellerie, F., 2011. Quelle unité pour le Châtelperronien? Apport de l’analyse taphonomique et techno-économique des industries lithiques de trois gisements aquitains de plein air: le Basté, Bidart (Pyrénées-Atlantiques) et Canaule II (Dordogne). Université Bordeaux 1, Bordeaux.

Bailey, S.E., Hublin, J.J., 2006a. Dental remains from the Grotte du Renne at Arcy-sur-Cure (Yonne). Journal of Human Evolution. 50, 485–508.

Bailey, S.E., Hublin, J.J., 2006b. Dental remains from the Grotte du Renne at Arcy-sur-Cure (Yonne). Journal of Human Evolution. 50, 485–508.

Baillet, M., Bachellerie, F., Bordes, J.-G., 2014. Investigation concerning a tool: techno-economical, functional and experimental analysis of Chatelperronian endscrapers from Canaule II (Creysse, Dordogne, France). Paléo. 07–36.

Bard, E., Heaton, T.J., Talamo, S., Kromer, B., Reimer, R.W., Reimer, P.J., 2020. Extended dilation of the radiocarbon time scale between 40,000 and 48,000 y BP and the overlap between Neanderthals and Homo sapiens. Proceedings of the National Academy of Sciences. 117, 21005–21007.

Barshay-Szmidt, C., Bazile, F., Brugal, J.-P., 2020. First AMS 14C dates on the Protoaurignacian in Mediterranean France: the site of Esquicho-Grapaou (Russan-Ste-Anastasie, Gard). Journal of Archaeological Science: Reports. 33, 102474.

Barshay-Szmidt, C., Normand, C., Flas, D., Soulier, M.C., 2018. Radiocarbon dating the Aurignacian sequence at Isturitz (France): Implications for the timing and development of the Protoaurignacian and Early Aurignacian in western Europe. Journal of Archaeological Science: Reports. 17, 809–838.

Barshay-Szmidt, C.C., Eizenberg, L., Deschamps, M., 2012. Radiocarbon (AMS) dating the Classic Aurignacian, Proto-Aurignacian and Vasconian Mousterian at Gatzarria Cave (Pyrénées-Atlantiques, France). PALEO. Revue d’archéologie préhistorique. 11–38.

Bar-Yosef, O., Bordes, J.G., 2010. Who were the makers of the Châtelperronian culture? Journal of Human Evolution. 59, 586–593.

Bergman, C., 1981. Point types in the Upper Palaeolithic sequence at Ksar’Akil, Lebanon. In: Cauvin, J., Sanlaville, P. (Eds.), Préhistoire Du Levant. C.N.R.S., Paris, pp. 319–330.

Bird, M.I., Beaman, R.J., Condie, S.A., Cooper, A., Ulm, S., Veth, P., 2018. Palaeogeography and voyage modeling indicates early human colonization of Australia was likely from Timor-Roti. Quaternary Science Reviews. 191, 431–439.

Bodu, P., Salomon, H., Lacarrière, J., Baillet, M., Balunger, M., Naton, H.G., Théry-Parisot, I., 2017. Un gisement châtelperronien de plein air dans le Bassin parisien: les Bossats à Ormesson (Seine-et-Marne). Gallia Prehistoire. 57, 3–64.

Boëda, E., 1991. Approche de la variabilité des systèmes de production lithique des industries du paléolithique inférieur et moyen: Chronique d’une variabilité attendue. Techniques et Culture. 17–18, 37–79.

Bordes, J.-G., 2003. Lithic taphonomy of the Châtelperronian/Aurignacian interstratifications of the Roc-de-Combe and Le Piage sites (Lot, france). In: Zilhão, J., D’Errico, F. (Eds.), The Chronology of the Aurignacian and of the Transitional Technocomplexes. Dating, Stratigraphies, Cultural Implications; Proceedings of Symposium 6.1 of the XIVth Congress of the UISPP (University of Liège, Belgium, September 2-8, 2001). IPA, Lisboa, pp. 223–244.

Bordes, J.-G., Teyssandier, N., 2011a. The Upper Paleolithic nature of the Châtelperronian in South-Western France: Archeostratigraphic and lithic evidence. Quaternary International. 246, 382–388.

Bordes, J.-G., Teyssandier, N., 2011b. The Upper Paleolithic nature of the Châtelperronian in South-Western France: Archeostratigraphic and lithic evidence. Quaternary International. 246, 382–388.

Bourne, M.D., Henderson, G.M., Thomas, A.L., Mac Niocaill, C., 2012. High-resolution palaeomagnetic records of the Laschamp geomagnetic excursion from ODP Sites 1061 and 1062. 2012, GP43A-1119.

Bricker, H.M., Laville, H., 1977. Le gisement châtelperronien de plein air des Tambourets (commune de Couladère, Haute-Garonne). Bulletin de la Société préhistorique française. Études et travaux. 74, 505–517.

Byrne, R.W., 2007. Culture in great apes: Using intricate complexity in feeding skills to trace the evolutionary origin of human technical prowess. In: Philosophical Transactions of the Royal Society B: Biological Sciences. Royal Society, pp. 577–585.

Caron, F., d’Errico, F., Del Moral, P., Santos, F., Zilhão, J., 2011. The reality of neandertal symbolic behavior at the grotte du renne, arcy-sur-cure, france. PLoS ONE. 6, e21545–e21545.

Clarkson, C., Jacobs, Z., Marwick, B., Fullagar, R., Wallis, L., Smith, M., Roberts, R.G., Hayes, E., Lowe, K., Carah, X., Florin, S.A., McNeil, J., Cox, D., Arnold, L.J., Hua, Q., Huntley, J., Brand, H.E.A., Manne, T., Fairbairn, A., Shulmeister, J., Lyle, L., Salinas, M., Page, M., Connell, K., Park, G., Norman, K., Murphy, T., Pardoe, C., 2017. Human occupation of northern Australia by 65,000 years ago. Nature. 547, 306–310.

Compton, T., Skinner, M.M., Humphrey, L., Pope, M., Bates, M., Davies, T.W., Parfitt, S.A., Plummer, W.P., Scott, B., Shaw, A., Stringer, C., 2021. The morphology of the Late Pleistocene hominin remains from the site of La Cotte de St Brelade, Jersey (Channel Islands). Journal of Human Evolution. 152, 102939.

Connet, N., 2002. Le Châtelperronien: Réflexions sur l’unité et l’identite techno-économique de l’industrie lithique. L’apport de l’analyse diachronique des industries lithiques des couches Châtelperronienes de la Grotte du Renne a Arcy-sur-Cure (Yonne). Universite de Lille 1.

d’Errico, F., Julien, M., Liolios, D., Vanhaeren, M., Baffier, D., 2003. Many Awls in Our Argument. Bone Tool Manufacture and Use in the Chatelperronian and Aurignacian Levels of the Grotte du Renne at Arcy-sur-Cure. In: Zilhão, J., d’Errico, F. (Eds.), The Chronology of the Aurignacian and of the Transitional Technocomplexes: Dating, Stratigraphies, Cultural Implications. Istituto Português de Arqueologia, Lisboa, pp. 247–270.

Demidenko, Y.E., Škrdla, P., 2023. Lincombian-Ranisian-Jerzmanowician Industry and South Moravian Sites: a Homo sapiens Late Initial Upper Paleolithic with Bohunician Industrial Generic Roots in Europe. Journal of Paleolithic Archaeology. 6, 17.

D’Errico, F., Zilhão, J., Julien, M., Baffler, D., Pelegrin, J., 1998. Neanderthal acculturation in western Europe? A critical review of the evidence and its interpretation. Current Anthropology. 39.

Devièse, T., Abrams, G., Hajdinjak, M., Pirson, S., Groote, I.D., Modica, K.D., Toussaint, M., Fischer, V., Comeskey, D., Spindler, L., Meyer, M., Semal, P., Higham, T., 2021. Reevaluating the timing of Neanderthal disappearance in Northwest Europe. Proceedings of the National Academy of Sciences. 118.

Discamps, E., 2011. Hommes et hyènes face aux recompositions des communautés d’Ongulés (MIS 5-3) : Éléments pour un cadre paléoécologique des sociétés du Paléolithique moyen et supérieur ancien d’Europe de l’Ouest, Bulletin de la Société préhistorique française. Université Sciences et Technologies - Bordeaux I.

Discamps, E., Bachellerie, F., Baoillet, M., Sitzia, L., 2019. The Use of Spatial Taphonomy for Interpreting Pleistocene Palimpsests: An Interdisciplinary Approach to the Châtelperronian and Carnivore Occupations at Cassenade (Dordogne, France). PaleoAnthropology. 362–388.

Djakovic, I., Key, A., Soressi, M., 2022. Optimal linear estimation models predict 1400–2900 years of overlap between Homo sapiens and Neandertals prior to their disappearance from France and northern Spain. Scientific Reports. 12, 15000.

Douka, K., Bergman, C.A., Hedges, R.E.M., Wesselingh, F.P., Higham, T.F.G., 2013. Chronology of Ksar Akil (Lebanon) and Implications for the Colonization of Europe by Anatomically Modern Humans. PLOS ONE. 8, e72931.

Essel, E., Zavala, E.I., Schulz-Kornas, E., Kozlikin, M.B., Fewlass, H., Vernot, B., Shunkov, M.V., Derevianko, A.P., Douka, K., Barnes, I., Soulier, M.-C., Schmidt, A., Szymanski, M., Tsanova, T., Sirakov, N., Endarova, E., McPherron, S.P., Hublin, J.-J., Kelso, J., Pääbo, S., Hajdinjak, M., Soressi, M., Meyer, M., 2023. Ancient human DNA recovered from a Palaeolithic pendant. Nature. 1–5.

Faivre, J.P., Discamps, E., Gravina, B., Turq, A., Bourguignon, L., 2017. Cleaning up a Messy Mousterian: How to describe and interpret Late Middle Palaeolithic chrono-cultural variability in Atlantic Europe. Quaternary International. 433, 1–3.

Falcucci, A., Conard, N.J., Peresani, M., 2017. A critical assessment of the Protoaurignacian lithic technology at Fumane Cave and its implications for the definition of the earliest Aurignacian, PLoS ONE.

Flas, D., 2011. The Middle to Upper Paleolithic transition in Northern Europe: The Lincombian-Ranisian-Jerzmanowician and the issue of acculturation of the last Neanderthals. World Archaeology. 43, 605–627.

Floss, H., Hoyer, C., Würschem, H., 2016. The Châtelperronian of Germolles (Grotte de la Verpillière I, Mellecey, Saône-et-Loire, France). Paléo. 149–176.

Gravina, B., Bachellerie, F., Caux, S., Discamps, E., Faivre, J.P., Galland, A., Michel, A., Teyssandier, N., Bordes, J.G., 2018. No Reliable Evidence for a Neanderthal-Châtelperronian Association at La Roche-à-Pierrot, Saint-Césaire. Scientific Reports. 8, 1–12.

Gravina, B., Discamps, E., 2015. MTA-B or not to be? Recycled bifaces and shifting hunting strategies at Le Moustier and their implication for the late Middle Palaeolithic in southwestern France. Journal of Human Evolution. 84, 83–98.

Gravina, B., Mellars, P., Ramsey, C.B., 2005. Radiocarbon dating of interstratified Neanderthal and early modern human occupations at the Chatelperronian type-site. Nature. 438, 51–56.

Grigoletto, F., Ortega, I., Rios, J., Bourguignon, L., 2008. Le Châtelperronien de Vieux Coutets. Premiers éléments de réflexion. In: Jaubert, J., Bordes, J.-G., Ortega, I. (Eds.), Les Sociétés Du Paléolithique Dans Un Grand Sud-Ouest: Nouveaux Gisements, Nouveaux Résultats, Nouvelles Méthodes. SPF, Paris, pp. 245–259.

Guilbaud, M., 1993. Debitage from the Upper Castelperronian level at Saint-Césaire: Methodological approach and implications for the transition from Middle to Upper Paleolithic. In: Léveque, F., Backer, A., Guilbaud, M. (Eds.), Context of a Late Neandertal. Prehistory Press, Madison, pp. 37–58.

Hajdinjak, M., Fu, Q., Hübner, A., Petr, M., Mafessoni, F., Grote, S., Skoglund, P., Narasimham, V., Rougier, H., Crevecoeur, I., Semal, P., Soressi, M., Talamo, S., Hublin, J.J., Gušić, I., Kućan, Z., Rudan, P., Golovanova, L.V., Doronichev, V.B., Posth, C., Krause, J., Korlević, P., Nagel, S., Nickel, B., Slatkin, M., Patterson, N., Reich, D., Prüfer, K., Meyer, M., Pääbo, S., Kelso, J., 2018. Reconstructing the genetic history of late Neanderthals. Nature. 555, 652–656.

Hajdinjak, M., Mafessoni, F., Skov, L., Vernot, B., Hübner, A., Fu, Q., Essel, E., Nagel, S., Nickel, B., Richter, J., Moldovan, O.T., Constantin, S., Endarova, E., Zahariev, N., Spasov, R., Welker, F., Smith, G.M., Sinet-Mathiot, V., Paskulin, L., Fewlass, H., Talamo, S., Rezek, Z., Sirakova, S., Sirakov, N., McPherron, S.P., Tsanova, T., Hublin, J.-J., Peter, B.M., Meyer, M., Skoglund, P., Kelso, J., Pääbo, S., 2021. Initial Upper Palaeolithic humans in Europe had recent Neanderthal ancestry. Nature. 592, 253–257.

Harrold, F.B., 2000. The Chatelperronian in historical context. Journal of Anthropological Research. 56, 59–75.

Harvati, K., Röding, C., Bosman, A.M., Karakostis, F.A., Grün, R., Stringer, C., Karkanas, P., Thompson, N.C., Koutoulidis, V., Moulopoulos, L.A., Gorgoulis, V.G., Kouloukoussa, M., 2019. Apidima Cave fossils provide earliest evidence of Homo sapiens in Eurasia. Nature.

Heaton, T.J., Blaauw, M., Blackwell, P.G., Ramsey, C.B., Reimer, P., Scott, E.M., 2020. The IntCal20 approach to radiocarbon calibration curve construction: A new methodology using Bayesian splines and errors-in-variables. Radiocarbon.

Higham, T., Douka, K., Wood, R., Ramsey, C.B., Brock, F., Basell, L., Camps, M., Arrizabalaga, A., Baena, J., Barroso-Ruíz, C., Bergman, C., Boitard, C., Boscato, P., Caparrós, M., Conard, N.J., Draily, C., Froment, A., Galván, B., Gambassini, P., Garcia-Moreno, A., Grimaldi, S., Haesaerts, P., Holt, B., Iriarte-Chiapusso, M.J., Jelinek, A., Jordá Pardo, J.F., Maíllo-Fernández, J.M., Marom, A., Maroto, J., Menéndez, M., Metz, L., Morin, E., Moroni, A., Negrino, F., Panagopoulou, E., Peresani, M., Pirson, S., De La Rasilla, M., Riel-Salvatore, J., Ronchitelli, A., Santamaria, D., Semal, P., Slimak, L., Soler, J., Soler, N., Villaluenga, A., Pinhasi, R., Jacobi, R., 2014. The timing and spatiotemporal patterning of Neanderthal disappearance. Nature. 512, 306–309.

Higham, T., Jacobi, R., Julien, M., David, F., Basell, L., Wood, R., Davies, W., Ramsey, C.B., 2010. Chronology of the Grotte du Renne (France) and implications for the context of ornaments and human remains within the Châtelperronian. Proceedings of the National Academy of Sciences of the United States of America. 107, 20234–20239.

Higham, T.F.G., Jacobi, R.M., Bronk Ramsay, C., 2006. AMS radiocarbon dating of ancient bone using ultrafiltration. Radiocarbon. 48, 179–195.

Hublin, J.-J., 2015. The modern human colonization of western Eurasia: when and where? Quaternary Science Reviews. 118, 194–210.

Hublin, J.J., Sirakov, N., Aldeias, V., Bailey, S., Bard, E., Delvigne, V., Endarova, E., Fagault, Y., Fewlass, H., Hajdinjak, M., Kromer, B., Krumov, I., Marreiros, J., Martisius, N.L., Paskulin, L., Sinet-Mathiot, V., Meyer, M., Pääbo, S., Popov, V., Rezek, Z., Sirakova, S., Skinner, M.M., Smith, G.M., Spasov, R., Talamo, S., Tuna, T., Wacker, L., Welker, F., Wilcke, A., Zahariev, N., McPherron, S.P., Tsanova, T., 2020. Initial Upper Palaeolithic Homo sapiens from Bacho Kiro Cave, Bulgaria. Nature. 581, 299–302.

Hublin, J.-J., Spoor, F., Braun, M., Zonneveld, F.W., Condemi, S., 1996. A late Neanderthal associated with Upper Palaeolithic aftefacts. Nature. 381, 224–226.

Hublin, J.J., Talamo, S., Julien, M., David, F., Connet, N., Bodu, P., Vandermeersch, B., Richards, M.P., 2012. Radiocarbon dates from the Grotte du Renne and Saint-Césaire support a Neandertal origin for the Châtelperronian. Proceedings of the National Academy of Sciences of the United States of America. 109, 18743–18748.

Jacobs, Z., Li, B., Jankowski, N., Soressi, M., 2015. Testing of a single grain OSL chronology across the Middle to Upper Palaeolithic transition at Les Cottés (France). Journal of Archaeological Science. 54, 110–122.

Jaubert, J., 2011. Les archéoséquences du Paléolithique moyen du Sud-Ouest de la France : quel bilan un quart de siècle après François Bordes ? In: Delpech, F., Jaubert, J. (Eds.), François Bordes et La Préhistoire. Colloque International François Bordes, Bordeaux 22-24 Avril 2009. Editions du CTHS, Paris, pp. 235–253.

Jaubert, J., Bordes, J.-G., Discamps, E., Gravina, B., 2011. A New Look at the End of the Middle Palaeolithic Sequence in Southwestern France. Actes du colloque “International Symposium Characteristic features of the Middle to Upper Paleolithic transition in Eurasia: development of culture and evolution of Homo species”, Denisova, 4-10 juillet 2011.

Kadowaki, S., Omori, T., Nishiaki, Y., 2015. Variability in Early Ahmarian lithic technology and its implications for the model of a Levantine origin of the Protoaurignacian. Journal of Human Evolution. 82, 67–87.

Kolobova, K.A., Roberts, R.G., Chabai, V.P., Jacobs, Z., Krajcarz, M.T., Shalagina, A.V., Krivoshapkin, A.I., Li, B., Uthmeier, T., Markin, S.V., Morley, M.W., O’Gorman, K., Rudaya, N.A., Talamo, S., Viola, B., Derevianko, A.P., 2020. Archaeological evidence for two separate dispersals of Neanderthals into southern Siberia. Proceedings of the National Academy of Sciences of the United States of America. 117, 2879–2885.

Kuhn, S.L., Stiner, M.C., Güleç, E., Özer, I., Yilmaz, H., Baykara, I., Açiklol, A., Goldberg, P., Martínez Molina, K., Ünay, E., Suata-Alpaslan, F., 2009. The early Upper Paleolithic occupations at Üçağızlı Cave (Hatay, Turkey). Journal of Human Evolution. 56, 87–113.

Leroi-Gourhan, A., 1958. Étude des restes humains fossiles provenant des Grottes d’Arcy-sur-Cure. Annales de Paléontologie. 44, 87–148.

Lévêque, F., Vandermeersch, B., 1981. Les restes humains de Saint-Cesaire (Charente-Maritime). Bulletins et Mémoires de la Société d’Anthropologie de Paris. 13–8, 103–104.

Marín-Arroyo, A.B., Rios-Garaizar, J., Straus, L.G., Jones, J.R., de la Rasilla, M., González Morales, M.R., Richards, M., Altuna, J., Mariezkurrena, K., Ocio, D., 2018. Chronological reassessment of the Middle to Upper Paleolithic transition and Early Upper Paleolithic cultures in Cantabrian Spain. PLoS ONE. 13, 1–20.

Massilani, D., Morley, M.W., Mentzer, S.M., Aldeias, V., Vernot, B., Miller, C., Stahlschmidt, M., Kozlikin, M.B., Shunkov, M.V., Derevianko, A.P., Conard, N.J., Wurz, S., Henshilwood, C.S., Vasquez, J., Essel, E., Nagel, S., Richter, J., Nickel, B., Roberts, R.G., Pääbo, S., Slon, V., Goldberg, P., Meyer, M., 2022. Microstratigraphic preservation of ancient faunal and hominin DNA in Pleistocene cave sediments. Proceedings of the National Academy of Sciences. 119, e2113666118.

Mellars, P., 2006. Archeology and the dispersal of modern humans in Europe: Deconstructing the “Aurignacian.” Evolutionary Anthropology. 15, 167–182.

Mellars, P., 2010. Neanderthal symbolism and ornament manufacture: The bursting of a bubble? Proceedings of the National Academy of Sciences of the United States of America. 107, 20147–20148.

Mellars, P., Gravina, B., Ramsey, C.B., 2007. Confirmation of Neanderthal/modern human interstratification at the Chatelperronian type-site. Proceedings of the National Academy of Sciences of the United States of America. 104, 3657–3662.

Metz, L., Lewis, J.E., Slimak, L., 2023. Bow-and-arrow, technology of the first modern humans in Europe 54,000 years ago at Mandrin, France. Science Advances. 9, eadd4675.

Norman, K., Inglis, J., Clarkson, C., Faith, J.T., Shulmeister, J., Harris, D., 2018. An early colonisation pathway into northwest Australia 70-60,000 years ago. Quaternary Science Reviews. 180, 229–239.

Oxilia, G., Bortolini, E., Marciani, G., Menghi Sartorio, J.C., Vazzana, A., Bettuzzi, M., Panetta, D., Arrighi, S., Badino, F., Figus, C., Lugli, F., Romandini, M., Silvestrini, S., Sorrentino, R., Moroni, A., Donadio, C., Morigi, M.P., Slon, V., Piperno, M., Talamo, S., Collina, C., Benazzi, S., 2022. Direct evidence that late Neanderthal occupation precedes a technological shift in southwestern Italy. American Journal of Biological Anthropology. 179, 18–30.

Pelegrin, J., 1995. Technologie lithique : le Châtelperronien de Roc-de-Combe (Lot) et de La Côte (Dordogne), Cahiers du Quaternaire. CNRS Editions, Paris.

Prüfer, K., Posth, C., Yu, H., Stoessel, A., Spyrou, M.A., Deviese, T., Mattonai, M., Ribechini, E., Higham, T., Velemínský, P., Brůžek, J., Krause, J., 2021. A genome sequence from a modern human skull over 45,000 years old from Zlatý kůň in Czechia. Nature Ecology & Evolution. 5, 820–825.

Rebollo, N.R., Weiner, S., Brock, F., Meignen, L., Goldberg, P., Belfer-Cohen, A., Bar-Yosef, O., Boaretto, E., 2011. New radiocarbon dating of the transition from the Middle to the Upper Paleolithic in Kebara Cave, Israel. Journal of Archaeological Science. 38, 2424–2433.

Rios-Garaizar, J., Iriarte, E., Arnold, L.J., Sánchez-Romero, L., Marín-Arroyo, A.B., Emeterio, A.S., Gómez-Olivencia, A., Pérez-Garrido, C., Demuro, M., Campaña, I., Bourguignon, L., Benito-Calvo, A., Iriarte, M.J., Aranburu, A., Arranz-Otaegi, A., Garate, D., Silva-Gago, M., Lahaye, C., Ortega, I., 2022. The intrusive nature of the Châtelperronian in the Iberian Peninsula. PLOS ONE. 17, e0265219.

Rocca, R., Connet, N., Lhomme, V., 2017. Avant la transitionLes industries du Paléolithique moyen final de la grotte du Renne (couche XI) à Arcy-sur-Cure (Bourgogne, France). Comptes Rendus - Palevol. 16, 878–893.

Roussel, M., 2011. Normes et variations de la production lithique durant le Châtelperronien : la séquence de la Grande-Roche-de-la-Plématrie à Quinçay (Vienne). Université Paris Ouest, Nanterre.

Roussel, M., Bourguignon, L., Soressi, M., 2009. Identification par l’expérimentation de la percussion au percuteur de calcaire au Paléolithique moyen: le cas du façonnage des racloirs bifaciaux Quina de Chez Pinaud (Jonzac, Charente-Maritime). Bulletin de la Société préhistorique française. 219–238.

Roussel, M., Soressi, M., 2006. Une nouvelle séquence du Paléolithique supérieur ancien aux marges sud-ouest du Bassin parisien : les Cottés dans la Vienne. Le Paléolithique supérieur ancien de l’Europe du Nord-ouest. Réflexions et synthèses à partir d’un projet collectif de recherche sur le centre et le sud du Bassin parisien, journées SPF, Sens, 15-18 avril 2009. Mémoire 56, Société préhistorique française. 283–297.

Roussel, M., Soressi, M., Hublin, J.J., 2016a. The Châtelperronian conundrum: Blade and bladelet lithic technologies from Quinçay, France. Journal of Human Evolution. 95, 13–32.

Roussel, M., Soressi, M., Hublin, J.J., 2016b. The Châtelperronian conundrum: Blade and bladelet lithic technologies from Quinçay, France. Journal of Human Evolution. 95, 13–32.

Ruebens, K., McPherron, S.J.P., Hublin, J.-J., 2015. On the local Mousterian origin of the Châtelperronian: Integrating typo-technological, chronostratigraphic and contextual data. Journal of Human Evolution. 86, 55–91.

Sier, M.J., Dekkers, M.J., Parés, J.M., Roebroeks, W., 2013. Land-sea correlation of the Last Interglacial via the Blake palaeomagnetic Event: implications for Neandertal occupation history of north western Europe. Proceedings of the European Society for the Study of Human Evolution. 2, 209–209.

Slimak, L., 2023. The three waves: Rethinking the structure of the first Upper Paleolithic in Western Eurasia. PLOS ONE. 18, e0277444.

Slimak Ludovic, Zanolli Clément, Higham Tom, Frouin Marine, Schwenninger Jean-Luc, Arnold Lee J., Demuro Martina, Douka Katerina, Mercier Norbert, Guérin Gilles, Valladas Hélène, Yvorra Pascale, Giraud Yves, Seguin-Orlando Andaine, Orlando Ludovic, Lewis Jason E., Muth Xavier, Camus Hubert, Vandevelde Ségolène, Buckley Mike, Mallol Carolina, Stringer Chris, Metz Laure, n.d. Modern human incursion into Neanderthal territories 54,000 years ago at Mandrin, France. Science Advances. 8, eabj9496.

Soressi, M., 2002. Le Moustérien de tradition acheuléenne du sud-ouest de la France. Discussion sur la signification du faciès à partir de l’étude comparée de quatre sites: Pech-de-l’Azé I, Le Moustier, La Rochette et la Grotte XVI. University of Bordeaux I, Bordeaux.

Soressi, M., 2011. Révision taphonomique et techno-typologique des deux ensembles attribués au Châtelperronien de la Roche-à-Pierrot à Saint-Césaire. L’Anthropologie, Paléolithique supérieur. 115, 569–584.

Soressi, M., Hublin, J.-J., Primault, J., Richards, M., Richte, r D., Rendu, W., Roussel, M., Texier, J.P., 2006. Les Cottés Saint-Pierre-de Maillé (Vienne) Rapport de fouille programmée 2006.

Soressi, M., McPherron, S.P., Lenoir, M., Dogandžić, T., Goldberg, P., Jacobs, Z., Maigrot, Y., Martisius, N.L., Miller, C.E., Rendu, W., Richards, M., Skinner, M.M., Steele, T.E., Talamo, S., Texier, J.-P., 2013. Neandertals made the first specialized bone tools in Europe. Proceedings of the National Academy of Sciences. 110, 14186–14190.

Soressi, M., Roussel, M., 2014. European Middle to Upper Palaeolithic Transitional Industries: Châtelperronian. In: Smith, Cl. (Ed.), Encyclopedia of Global Archaeology. Springer, New York, pp. 2679–2693.

Soriano, S., Villa, P., Wadley, L., 2007. Blade technology and tool forms in the Middle Stone Age of South Africa: the Howiesons Poort and post-Howiesons Poort at Rose Cottage Cave. Journal of Archaeological Science. 34, 681–703.

Stringer, C., Crété, L., 2022. Mapping Interactions of H. neanderthalensis and Homo sapiens from the Fossil and Genetic Records. PaleoAnthropology. 2022.

Talamo, S., Aldeias, V., Goldberg, P., Chiotti, L., Dibble, H.L., Guérin, G., Hublin, J.-J., Madelaine, S., Maria, R., Sandgathe, D., Steele, T.E., Turq, A., Mcpherron, S.J.P., 2020. The new 14C chronology for the Palaeolithic site of La Ferrassie, France: the disappearance of Neanderthals and the arrival of Homo sapiens in France. Journal of Quaternary Science. 35, 961–973.

Talamo, S., Soressi, M., Roussel, M., Richards, M., Hublin, J.J., 2012. A radiocarbon chronology for the complete Middle to Upper Palaeolithic transitional sequence of Les Cottés (France). Journal of Archaeological Science. 39, 175–183.

Teyssandier, N., Bon, F., Bordes, J.G., 2010. Within projectile range: Some thoughts on the appearance of the Aurignacian in Europe. Journal of Anthropological Research. 66, 209–229.

Tobler, R., Rohrlach, A., Soubrier, J., Bover, P., Llamas, B., Tuke, J., Bean, N., Abdullah-Highfold, A., Agius, S., O’Donoghue, A., O’Loughlin, I., Sutton, P., Zilio, F., Walshe, K., Williams, A.N., Turney, C.S.M., Williams, M., Richards, S.M., Mitchell, R.J., Kowal, E., Stephen, J.R., Williams, L., Haak, W., Cooper, A., 2017. Aboriginal mitogenomes reveal 50,000 years of regionalism in Australia. Nature. 544, 180–184.

Vallini, L., Marciani, G., Aneli, S., Bortolini, E., Benazzi, S., Pievani, T., Pagani, L., 2022. Genetics and material culture support repeated expansions into Paleolithic Eurasia from a population Hub out of Africa. Genome Biology and Evolution. evac045.

Welker, F., Hajdinjak, M., Talamo, S., Jaouen, K., Dannemann, M., David, F., Julien, M., Meyer, M., Kelso, J., Barnes, I., Brace, S., Kamminga, P., Fischer, R., Kessler, B.M., Stewart, J.R., Pääbo, S., Collins, M.J., Hublin, J.J., 2016. Palaeoproteomic evidence identifies archaic hominins associated with the Châtelperronian at the Grotte du Renne. Proceedings of the National Academy of Sciences of the United States of America. 113, 11162–11167.

Welker, F., Soressi, M.A., Roussel, M., van Riemsdijk, I., Hublin, J.J., Collins, M.J., 2017. Variations in glutamine deamidation for a Châtelperronian bone assemblage as measured by peptide mass fingerprinting of collagen. Science and Technology of Archaeological Research. 3, 15– 27.

Wiśniewski, A., Pyżewicz, K., Serwatka, K., Kot, M., Kerneder-Gubała, K., Grużdź, W., 2022. Lincombian-Ranisian-Jerzmanowician points were used primarily as hunting weapons: morphological and functional analysis of points from Nietoperzowa Cave, southern Poland. Archaeological and Anthropological Sciences. 14, 90.

Ziffer, D., 1978. A Re-Evaluation of the Upper Palaeolithic Industries at Kebara Cave and Their Place in the Aurignacian Culture of the Levant. Paléorient. 4, 273–293.

Zwyns, N., 2012. Laminar technology and the onset of the Upper Paleolithic in the Altai, Siberia.

Zwyns, N., 2021. The Initial Upper Paleolithic in Central and East Asia: Blade Technology, Cultural Transmission, and Implications for Human Dispersals. Journal of Paleolithic Archaeology. 4, 19.

